# Force transmission balance through adhesions determines multicellular handedness

**DOI:** 10.64898/2026.04.04.716462

**Authors:** Tomoki Ishibashi, Ryohei Nishizawa, Goshi Ogita, Naoko Tokushige, Tatsuo Shibata

## Abstract

Cell chirality has been implicated in left-right (LR) asymmetric morphogenesis, yet multicellular handedness is not a simple readout of single-cell chirality. Here, we show that multicellular LR asymmetry is determined by the balance of force transmission through cell-cell and cell-substrate adhesions. Combining experiments in human epithelial cells with theoretical modeling, we demonstrate that clockwise-rotating single cells can generate either clockwise or counterclockwise collective rotation depending on whether chiral forces are transmitted primarily through adherens junctions or focal adhesions, without altering single-cell chirality itself. Mechanistically, microtubules promoted junctional force transmission by recruiting the actin-microtubule crosslinker ACF7 to cell-cell contacts, where ACF7 stabilized F-actin and enabled intercellular mechanical coupling. Our results define a mechanical framework in which the routing of chiral forces through adhesions, rather than single-cell chirality alone, determines multicellular handedness.

## Main

Chirality, or left-right (LR) asymmetry, is a fundamental property of living systems and spans biological scales from molecules (e.g., L-amino acids) to organs (e.g., organ morphology). Macroscopic chirality at the organismal scale is thought to derive from molecular chirality, as organisms on Earth are built from homochiral components^1,2^. However, the handedness of macroscopic chirality can sometimes reverse, as in the development of a mirrored heart, suggesting that it does not arise through a simple assembly of microscopic chirality but instead emerges as a collective property of interacting chiral elements. Given that macroscopic handedness is not a direct reflection of the chirality of its homochiral elements, how do homochiral systems generate LR asymmetry while retaining the capacity for reversal?

Recent studies have proposed cell chirality as an intermediate scale linking molecules to tissues^3,4^. Cells display intrinsic LR asymmetry arising from cytoskeletal mechanics, particularly actomyosin-generated chiral torque^2,5–10^. Because cells generate cytoskeletal forces and transmit them to neighboring cells and the substrate, they act as mechanical intermediates that couple molecular chirality to multicellular chirality. Multicellular chirality is therefore an emergent property arising from force transmission within and between cells. The central unresolved questions are how actomyosin-dependent chiral forces are coupled to adhesion systems to enable intracellular force relay, and how these forces are directed through cell-cell versus cell-substrate interfaces.

### Chiral collective rotation in Caco-2 cells

To address these cross-scale questions, we used the human colon cancer cell line, Caco-2, as a model system. Previously, we showed that isolated Caco-2 cells exhibit intrinsic cell chirality^10^. In singly isolated Caco-2 cells, the nucleus and cytoplasm rotated clockwise at an average speed of 50 degrees per hour when viewed from the apical side (Extended Data Fig. 1a,b; Supplementary Video 1)^10^. As Caco-2 cells spontaneously organize into epithelial colonies, we examined the collective behavior of these epithelial colonies (Supplementary Video 2).

Interestingly, the colonies exhibited clockwise rotation (Extended Data Fig. 2a; Supplementary Video 2). Although clockwise cell migration was transiently disturbed immediately before and after cell division, cells soon resumed stable clockwise movement (Extended Data Fig. 2b). The colonies appeared to undergo coherent rotation around a single rotation center near the colony center, rather than around multiple rotational axes, at least up to about 20 cells. This chiral collective behavior was already evident in two-cell colonies, which exhibited the highest angular velocity (Extended Data Fig. 2c). To understand the fundamental principles underlying collective rotation arising from cell chirality, we focused on two-cell colonies as the simplest multicellular system (Fig. 1a; Supplementary Video 3).

**Figure 1:**
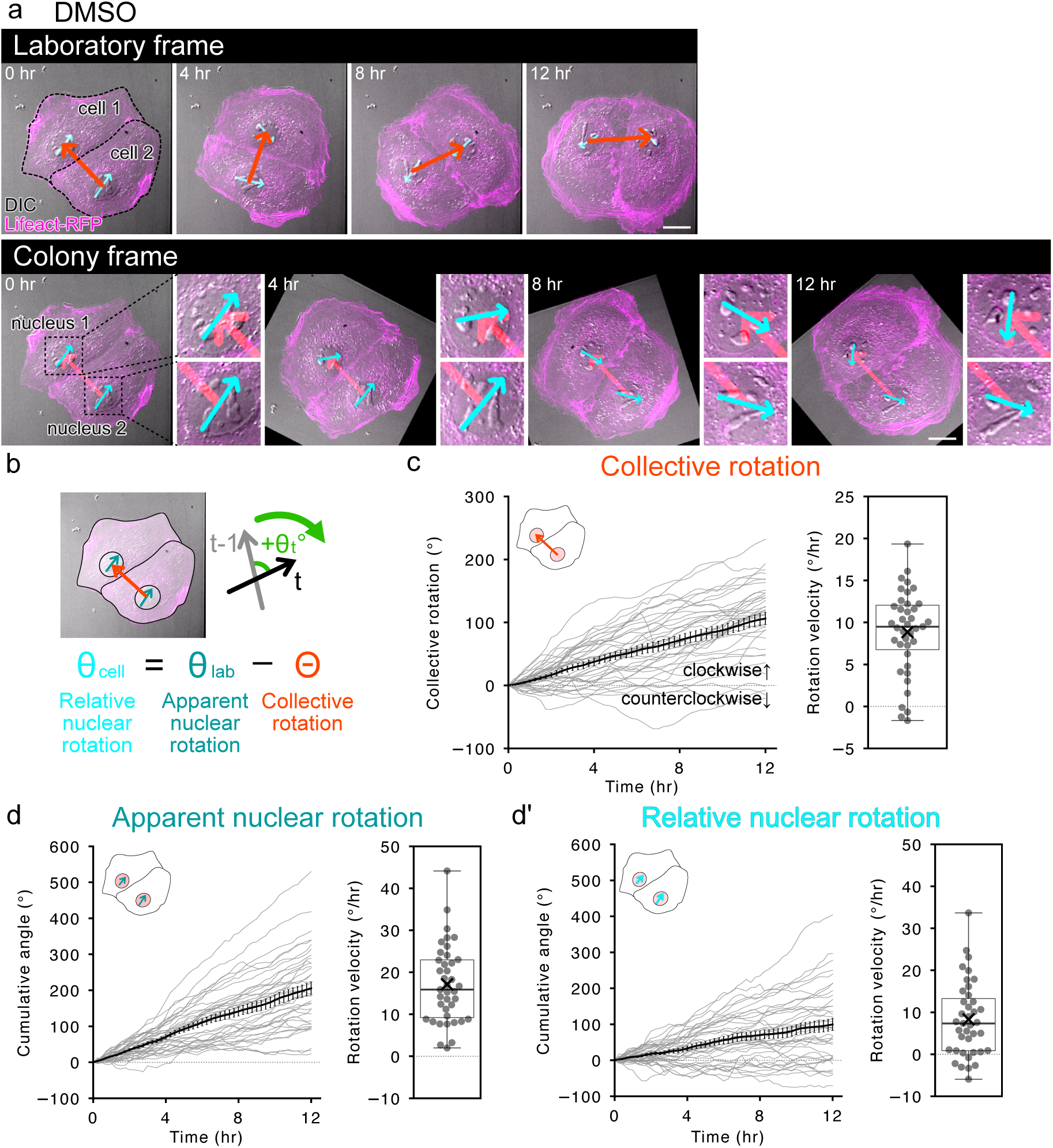
Caco-2 cells collectively rotate clockwise. **a**, Representative time course of a paired Caco-2 colony showing clockwise collective rotation. Time-lapse images of DIC (gray) and Lifeact-RFP (magenta) are shown. Images in the laboratory frame (top) and colony frame (bottom) are shown. Cell outlines are shown as dashed lines in the top panel. Cyan arrows are the nuclear vectors connecting two nuclear speckles, which were used to measure nuclear rotation. Red arrows are the vectors connecting the midpoints of the nuclear vectors, which were used to measure collective rotation. Note that the angles of the red arrows do not change in the colony frame images (bottom). In the colony frame images, enlarged views of the two cell nuclei are shown (bottom). Scale bars: 20 *μ*m. **b**, Schematic of the procedure used to quantify rotation. In laboratory frame images, two distinct nuclear speckles were selected, and the angle of the vector (dark blue) connecting them was measured to quantify apparent nuclear rotation (*θ_lab_*). The midpoints of the nuclear vectors were used as nuclear positions, and the angle of the vector (orange) connecting the nuclear positions was measured to quantify collective rotation (*Θ*). The angle between the vectors in the current (*t*) and previous (t − 1) frames was measured. Relative nuclear rotation (nuclear rotation in the colony frame; *θ_cell_*) was calculated by subtracting collective rotation from apparent nuclear rotation. **c-d**, Quantification of collective rotation (c), apparent nuclear rotation (d), and relative nuclear rotation (d’) in control cells (DMSO-treated). Left panels: Time courses of cumulative rotation angles over 12 h. Black solid lines indicate mean cumulative angles, and gray lines indicate cumulative angles of individual colonies or cells. Error bars indicate standard errors. Hereafter, positive angle values indicate clockwise rotation. Right panels: Box plots of angular velocities calculated for 12 h of observation. Circles indicate angular velocities of individual colonies or cells. Crosses indicate mean values. Sample size, *n* = 39.

The collective rotation speed (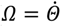; *Θ* is defined in Fig. 1b) averaged about ten degrees per hour in the clockwise direction (Fig. 1c). Approximately 90% of colonies exhibited clockwise collective rotation, whereas the counterclockwise-rotating colonies showed lower rotational speeds (Fig. 1c). The apparent angular velocity of nuclei in the laboratory frame (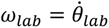; Fig. 1b) was approximately 20 degrees per hour in the clockwise direction (Fig. 1d). To estimate intrinsic cell rotation, we calculated the relative angular velocity of the nucleus in the co-rotating frame of the colony as *ω_cell_* = *ω_lab_* − *Ω* (Fig. 1a, bottom; Fig. 1b; Supplementary Video 3). Importantly, nuclear rotation remained clockwise even in the rotating colony frame, indicating that the intracellular mechanism generating rotational force remains active in two-cell colonies. However, the average relative nuclear rotation speed was approximately ten degrees per hour (Fig. 1d’), which was fivefold lower than that of a single-cell (Extended Data Fig. 1b). This reduction suggests that part of the rotational force generated by individual cells is transmitted through cell-cell interactions to drive the rotation of the colony as a whole. Hereafter, we used this relative angular velocity as an index of individual cell chirality during collective rotation.

### Distinct roles of actomyosin and microtubules in collective rotation

To investigate the forces driving collective rotation and relative nuclear rotation, we next assessed the contributions of the actomyosin and microtubule cytoskeletons. In two-cell colonies, F-actin was detected at the cell periphery as well as at cell-cell contact sites along the junction (Fig. 2a, top panel). Microtubules exhibited a sinistral chiral pattern, as observed in singly isolated cells^10^, and were broadly distributed throughout the cytoplasm (Fig. 2a, middle panel).

**Figure 2:**
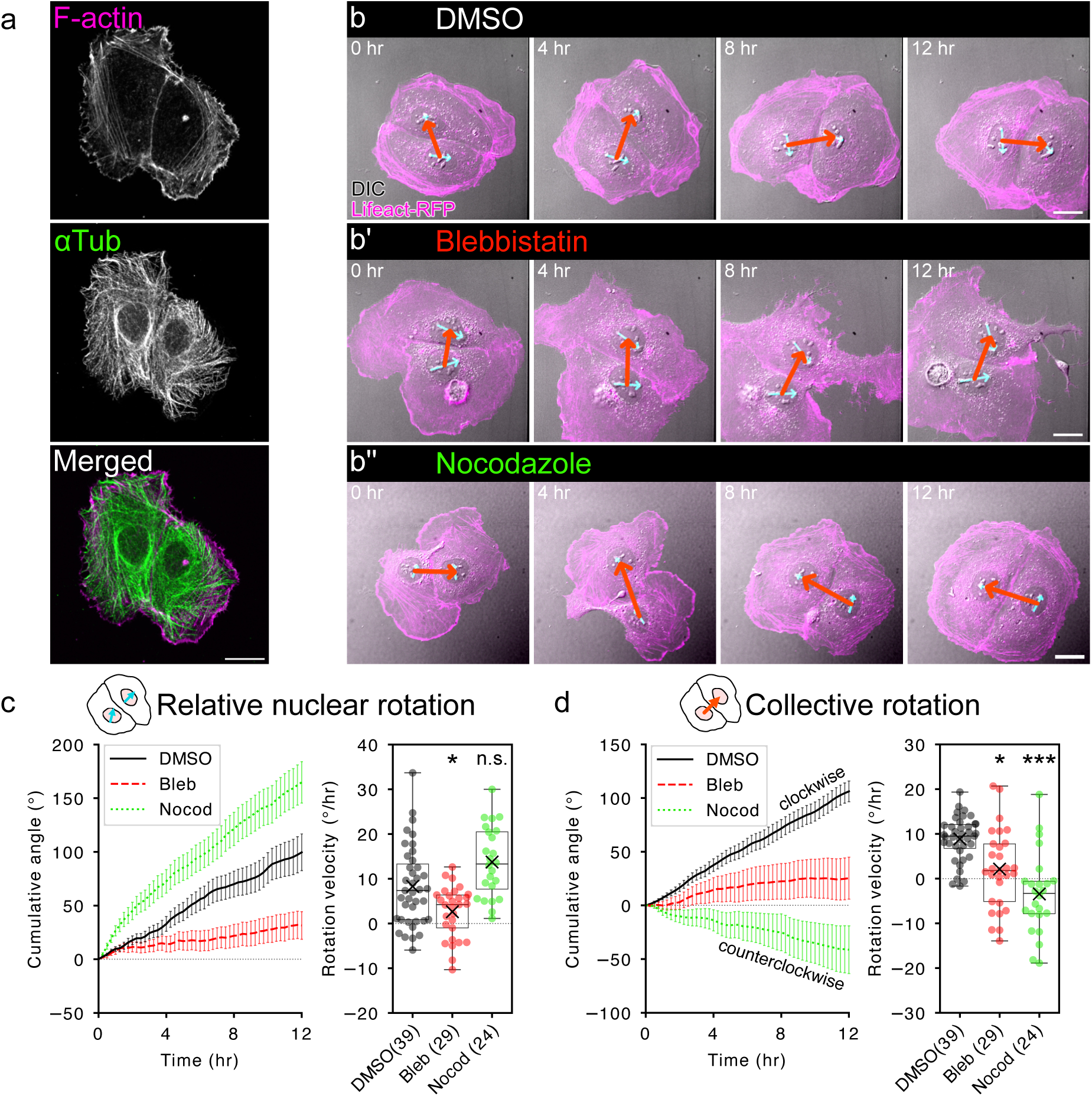
Contribution of actomyosin and microtubule cytoskeletons to collective rotation. **a**, Cytoskeletal organization of F-actin (top, phalloidin, magenta), microtubule (middle, anti-*α* tubulin, green), and merged image (bottom) are shown. **b**, Representative time courses of two-cell colonies treated with DMSO (b), blebbistatin (b’), or nocodazole (b’’) are shown in the laboratory frame. Time-lapse images of DIC (gray) and Lifeact-RFP (magenta) are shown. Cyan and red arrows are the vectors used to measure nuclear rotation and collective rotation, respectively. Scale bars: 20 *μ*m. **c-d**, Quantification of relative nuclear rotation (c) and collective rotation (d). Left panels: Time courses of cumulative rotation angles over 12 h. Black solid, red dashed, and green dotted lines indicate cumulative rotation angles in control (DMSO-treated), blebbistatin-treated, and nocodazole-treated cells, respectively. Error bars indicate standard errors. Right panels: Box plots of angular velocities calculated for 12 h of observation. Circles indicate angular velocities of individual cells. Crosses indicate mean values. Sample sizes are shown in parentheses. n.s.: *p* > 0.05; *: *p* < 0.05; ***: *p* < 0.001 by *t*-test followed by multiple pairwise comparisons with Bonferroni’s correction.

To test whether actomyosin, microtubules, or both contribute to collective rotation, we inhibited myosin II and microtubule polymerization using blebbistatin and nocodazole, respectively. Blebbistatin treatment inhibited relative nuclear rotation in two-cell colonies (Fig. 2b,b’,c; Supplementary Video 4), similar to its effect on single-cell rotation (Extended Data Fig. 1b; Supplementary Video 5). In addition, it randomized the direction of collective rotation while significantly reducing the rotation speed (Fig. 2d; Supplementary Video 4). These results indicate that the actomyosin cytoskeleton is required for both single-cell and multicellular chirality and suggest that actomyosin-dependent forces drive collective rotation. We next examined the role of microtubules by disrupting microtubule polymerization with nocodazole (Fig. 2b’’; Supplementary Videos 6 and 7). Nocodazole-treated colonies showed a slight increase in relative nuclear rotation, but the difference was not statistically significant (Fig. 2c). Surprisingly, approximately 80% of colonies rotated counterclockwise even though the nuclei continued to rotate clockwise, with an average speed of about five degrees per hour (Fig. 2d; Supplementary Video 7). These results suggest that microtubules regulate handedness at the multicellular level.

### Determination of collective rotation direction by the balance of force transmission through cell-cell and cell-substrate adhesions

To explain the reversal of collective rotation without reversal of individual cell chirality, we focused on force transmission and noted an analogous behavior in mechanical engineering. In automobiles, both forward and reverse motion can be generated from the same direction of engine rotation through a gear mechanism known as a planetary gear train, which consists of a sun gear, planetary gears, and an outer ring gear (Supplementary Figure 1a). In this system, when a clockwise-spinning planetary gear is coupled to the sun gear, it revolves clockwise around the sun gear (Supplementary Figure 1b). By contrast, when the planetary gear is coupled to the outer ring gear, it revolves counterclockwise even though it continues to spin clockwise (Supplementary Figure 1c). We therefore considered the possibility that, in a two-cell colony, the two cells behave analogously to the planetary and sun gears, with cell-cell adhesion corresponding to planetary-sun coupling and cell-substrate adhesion corresponding to planetary-outer ring coupling.

To test this idea, we developed a theoretical description of collective rotation using a two-dimensional cell membrane model (see Supplementary Notes for details). In the model, cell membrane and actin cortex are represented by connected segments, such that each cell is described as a polygon with multiple vertices (Fig. 3a). To simulate chiral cell rotation, we introduced a clockwise tangential active force at individual vertices (Fig. 3a), cell-cell adhesion between vertices at the interfaces between the two cells, and frictional forces at individual vertices to represent cell-substrate adhesion (Fig. 3b). Because focal adhesions (FAs) are not well developed near the cell-cell boundary in our Caco-2 cells (Supplementary Figure 1c), the frictional force against the substrate was set to be higher along the colony periphery than along the cell-cell interface.

**Figure 3:**
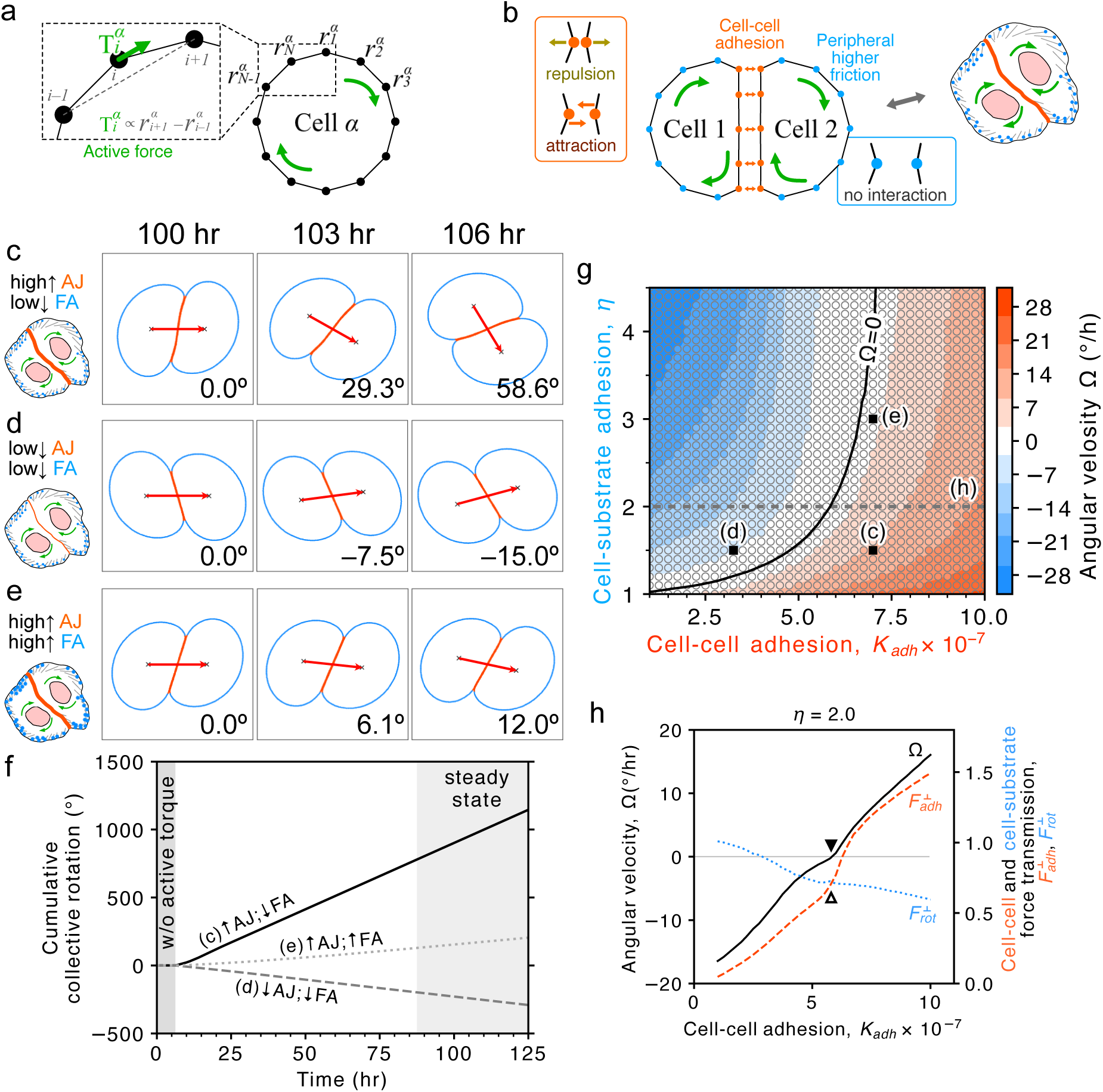
Theoretical simulation of collective rotation in Caco-2 cells. **a-b**, Schematic of the theoretical model used to reproduce collective rotation in a two-cell Caco-2 colony. Clockwise tangential active forces 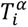 were applied to each cell vertex (a). A two-cell colony was formed through cell-cell adhesion (b, orange vertices). To represent cell-substrate adhesion, higher friction was assigned to vertices lacking cell-cell contacts (b, blue vertices). **c-e**, Simulation results under different levels of cell-cell and cell-substrate adhesion. Under relatively strong cell-cell adhesion and weak cell-substrate adhesion, cells collectively rotated clockwise (c). Under weaker cell-cell adhesion and similarly weak cell-substrate adhesion, cells collectively rotated counterclockwise (d). Under strong cell-cell adhesion and strong cell-substrate adhesion, slow clockwise collective rotation was observed (e). Red arrows connect the centroids of the two cells. **f**, Quantification of collective rotation. The conditions shown in c, d, and e correspond to the solid, dashed, and dotted lines, respectively. **g**, Phase diagram of collective rotation angular velocity (*Ω*) on the *K_adh_* − *η* plane. The color scale denotes the value of *Ω*. Circles indicate the parameter sets used, and their colors indicate the corresponding simulated values of *Ω*. The background was interpolated from the simulation data. The black line indicates the estimated contour at *Ω* = 0. Black squares indicate the parameter sets used in c, d, and e. **h**, Relationship between *Ω* and cell-cell or cell-substrate force transmission (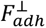 and 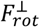, respectively). The *x*-axis indicates the cell-cell adhesion parameter *K_adh_*). For cell-substrate adhesion, *η* = 2.0 was used. *Ω*, 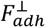, and 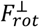 are shown as black solid, orange dashed, and blue dotted lines, respectively. Note that when the relative magnitude of 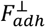 and 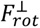 is reversed (white triangle), collective rotation *Ω* switches from a negative to a positive value (black triangle).

We next asked whether numerical simulation could recapitulate the *in cellulo* collective rotation (Fig. 3c-f). To simulate the control condition, we used relatively strong cell-cell adhesion to represent strong mechanical coupling between the two cells, relatively low peripheral friction, and a clockwise active force. Under this condition, the model reproduced clockwise collective rotation together with relative clockwise rotation of individual cells (Fig. 3c,f; Supplementary Video 8).

To test the effect of reduced cell-cell coupling, we decreased the cell-cell adhesion parameter without changing the cell-substrate frictional force and found that the colony rotated counterclockwise (Fig. 3d,f; Supplementary Video 9). We also tested the contribution of cell-substrate adhesion to collective rotation direction. When cell-substrate friction was increased without changing the cell-cell adhesion parameter from the control condition, the clockwise-rotating cells exhibited slower clockwise collective rotation than in the control (Fig. 3e,f; Supplementary Video 10). Furthermore, increasing cell-substrate friction also decreased the rotational speed at the single-cell level.

We then systematically investigated how collective rotation speed, *Ω*, depends on the balance between the cell-cell and cell-substrate adhesion, whose strengths are parametrized by *K_adh_* and *η*, respectively (see Supplementary Notes for details). As *K_adh_* decreased, *Ω* continuously decreased to zero and eventually became negative, indicating counterclockwise collective rotation (Fig. 3g). Similarly, as *η* increased, *Ω* continuously decreased and could also reach negative values corresponding to counterclockwise collective rotation (Fig. 3g). In summary, these results indicate that collective rotation can arise from individual cell chirality, whereas the direction of collective rotation is dictated by the balance between cell-cell and cell-substrate adhesion strengths.

To further examine how changes in mechanical interactions between the two cells reverse collective rotation direction, we measured the force exerted by one cell on the other and how this force changes with adhesion strength (see Methods for details). We found that only the tangential component of the cell-cell adhesion force (i.e., the component perpendicular to the line connecting the two cell centroids) contributes to collective rotation (see Eq. (2) in Methods). As *K_adh_* decreased, the tangential component of the force exerted by one cell on the other also decreased, indicating reduced force transmission between two cells (orange dashed line in Fig. 3h). Furthermore, when the direction of collective rotation reversed (black triangle in Fig. 3h), the relative magnitudes of the tangential cell-cell adhesion force (orange dashed line in Fig. 3h) and the cell-substrate frictional force arising from individual cell rotation were also reversed (white triangle in Fig. 3h; see also Eq. (2) in Methods). These results show that force transmission changes as a function of cell-cell adhesion strength. Thus, the balance of force transmission through cell-cell and cell-substrate adhesions determines the direction of collective rotation.

### Regulation of collective rotation direction by adherens junctions and focal adhesions

Our theory predicts that the relative strength of cell-cell and cell-substrate adhesion determines the direction of collective rotation (Fig. 4a). To test our hypothesis, we manipulated the balance between cell-cell and cell-substrate force transmission by inhibiting the core component of adherens junctions (AJs) using an inhibitory antibody against E-Cadherin (ECad). SHE78-7 is an inhibitory antibody against ECad, and it strongly suppresses ECad-mediated cell aggregation by binding to the EC1 domain of ECad^11^. First, we confirmed that neither SHE78-7 nor control IgG treatment affected single-cell rotation (Fig. 4b-d; Extended Data Fig. 3a,b; Supplementary Videos 11 and 12), indicating that ECad-dependent cell-cell adhesion is not required for cell chirality. Next, we observed the collective rotation in SHE78-7 treated colonies (Fig. 4e) and found that approximately two-thirds of them exhibited counterclockwise collective rotation, with an average speed of about five degrees per hour. In SHE78-7 treated colonies, cell pairs frequently separated and reattached, indicating decreased cell-cell adhesion (Supplementary Videos 13 and 14). These results indicate that the strength of cell-cell mechanical coupling determines the direction of collective rotation.

**Figure 4:**
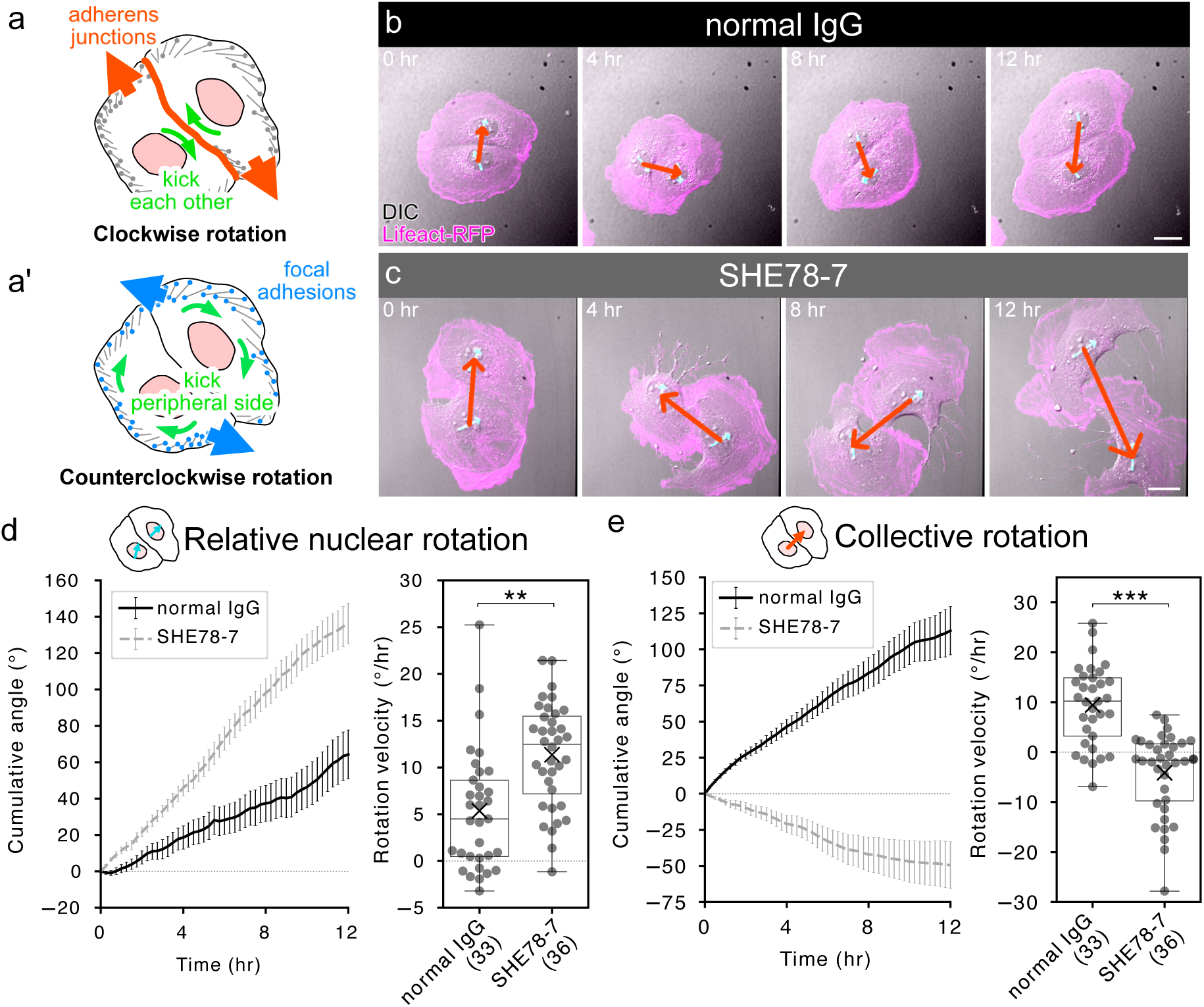
Inhibition of adherens junction reverses collective rotation. **a**, Schematic of our hypothesis that the balance of cell-cell adhesions and cell-substrate adhesions dictates the direction of collective rotation. When the relative contribution of cell-cell adhesions (e.g., adherens junctions, orange) is greater than that of cell-substrate adhesions (e.g., focal adhesions, blue), cells collectively rotate clockwise by kicking each other (a). When the relative contribution of cell-substrate adhesions is greater, cells collectively rotate counterclockwise by kicking the colony periphery (a’). **b-c**, Representative time courses of paired Caco-2 cells treated with normal IgG (b, control) and SHE78-7 (c). Time-lapse images of DIC (gray) and Lifeact (magenta) are shown. Cyan and red arrows are the vectors used to measure nuclear rotation and collective rotation, respectively. Scale bars: 20 *μ*m. **d-e**, Quantification of collective rotation (d) and relative nuclear rotation (e). Left panels: Time courses of cumulative rotation over 12 h. Black and gray lines indicate cumulative rotation angles in normal IgG treated and SHE78-7 treated cells, respectively. Error bars indicate standard errors. Right panels: Box plots of angular velocities calculated for 12 h of observation. Circles indicate angular velocities of individual cells. Crosses indicate average values. Sample sizes are shown in parentheses. **: *p* < 0.01; ***: *p* < 0.001 by *t*-test.

We next perturbed cell-substrate force transmission by overexpressing focal adhesion kinase (FAK) (Extended Data Fig. 4). FAK is involved in the assembly and maturation of FAs^12^. In cells overexpressing GFP-tagged FAK (GFP::FAK), the amount of activated FAK (pFAK; FAK pTyr397) increased, suggesting that FAK overexpression enhanced cell-substrate adhesion (Extended Data Fig. 4a). We then examined the effect of GFP::FAK overexpression on collective rotation, and found that it suppressed clockwise collective rotation (Extended Data Fig. 4b,c; Supplementary Videos 15 and 16). Notably, GFP::FAK overexpression did not alter relative nuclear rotation (Extended Data Fig. 4d), suggesting that the cells within the colony still generated sufficient chiral force, although GFP::FAK overexpression significantly attenuated single-cell rotation compared with GFP alone (Extended Data Fig. 4e,f; Supplementary Videos 17 and 18). These results suggest that enhanced cell-substrate adhesion counteracts the transmission of rotational forces through cell-cell interfaces, thereby weakening the clockwise collective rotation driven by cell chirality. Overall, our findings support the idea that the mechanical balance between cell-cell and cell-substrate adhesions governs the direction of collective rotation independently of the intrinsic chirality of individual cells.

### Regulation of intercellular and intracellular force transmission by microtubules and ACF7

Given that the microtubule disruption induced reversal of collective rotation in nocodazole-treated colonies (Fig. 2b’’,d), we hypothesized that microtubule disruption alters the balance of force transmission between cell-cell and cell-substrate adhesions, thereby reversing collective rotation. Previous studies have shown that microtubule disruption promotes FA activation^13^; therefore, we first examined cell-substrate adhesion. To assess FA formation, we stained for activated FAK (pFAK). In both control and nocodazole-treated colonies, pFAK signal was detected around the colony periphery, but not in the colony center (Fig. 5a). Compared with control colonies, nocodazole-treated colonies exhibited increased numbers and larger sizes of pFAK-positive speckles (Fig. 5b), indicating that microtubule disruption enhances cell-substrate adhesion.

**Figure 5:**
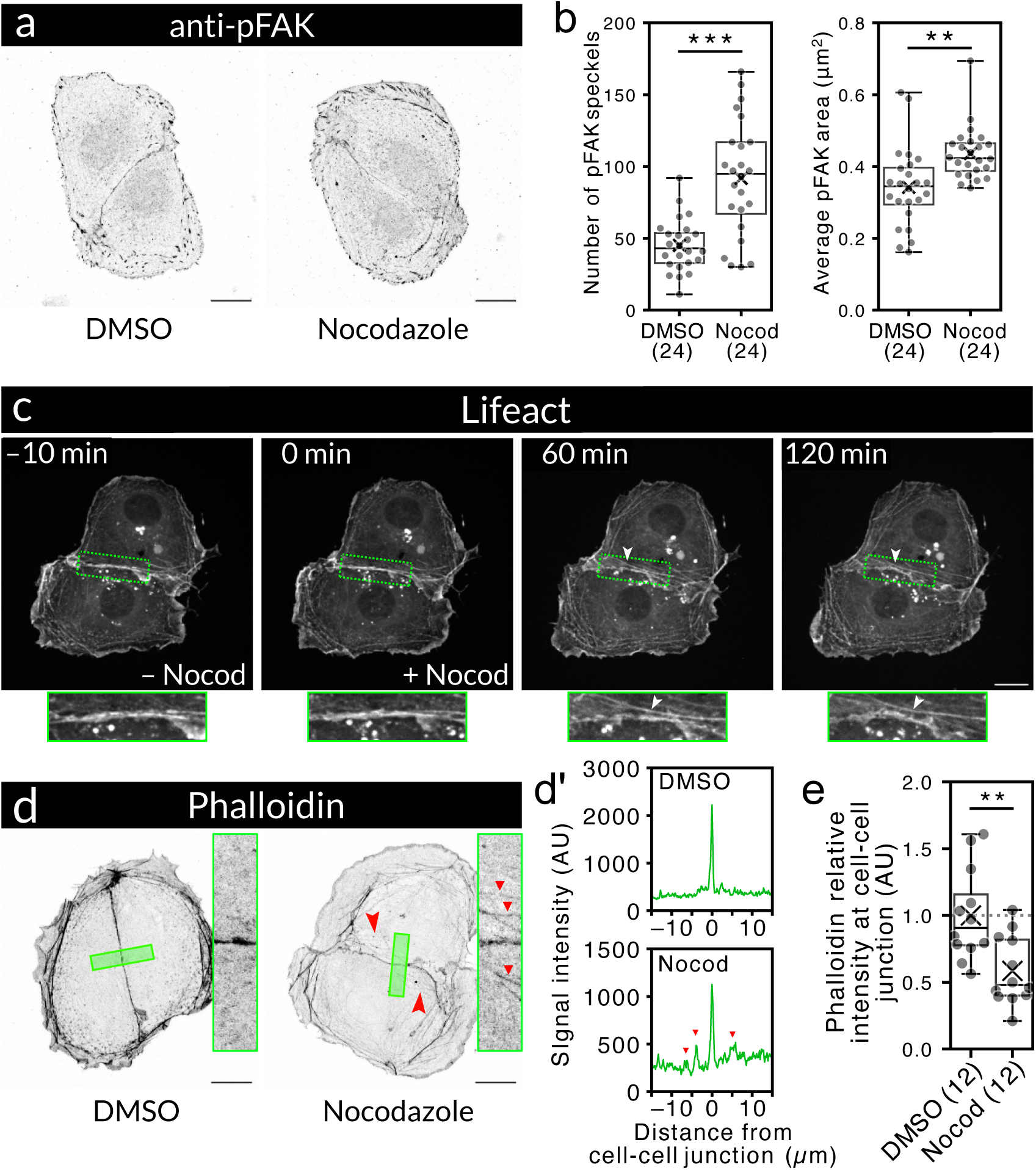
Microtubule disruption increases focal adhesions and decreases F-actin at cell-cell junctions. **a**, Representative images of anti-pFAK staining in DMSO-treated (left) and nocodazole-treated cells (right). **b**, Box plots of the number of pFAK speckles (left) and their average area (right). **c**, Representative time-lapse images of Lifeact-RFP immediately before (−10 min) and after nocodazole addition (0, 60, and 120 min) are shown. Regions outlined by green rectangles near the cell-cell boundary are shown at higher magnification in the bottom panels. White arrowheads indicate F-actin detached from the cell-cell boundary. **d**, Representative images of phalloidin staining of DMSO-treated (left) and nocodazole-treated cells (right). Regions outlined by green rectangles near the cell-cell boundary are shown at higher magnification in the right panels, and phalloidin signal intensity across these regions is plotted as a function of distance from the cell-cell boundary in d’. Red arrowheads and triangles indicate F-actin detached from the cell-cell boundary. **e**, Box plot of relative phalloidin intensity at the cell-cell boundary. Scale bars: 20 *μ*m. **: *p* < 0.01; ***: *p* < 0.001 by Mann-Whitney’s *U* test.

Next, to evaluate the effect of microtubule disruption on intercellular force transmission, we examined AJ. We obtained immunofluorescence images of ECad (Extended Data Fig. 5a) and quantified AJ levels by measuring ECad fluorescence intensity along the cell-cell boundary (Extended Data Fig. 5b,c). In nocodazole-treated colonies, ECad localization at the cell-cell junction was slightly decreased compared with that in control colonies, but the difference was not statistically significant (Extended Data Fig. 5c). Interestingly, however, F-actin frequently detached from cell-cell junctions after nocodazole treatment (14 of 16 colonies; Fig. 5c; Supplementary Videos 19; red arrowheads and triangles in Fig. 5d,d’). Furthermore, nocodazole treatment significantly decreased F-actin localization along cell-cell junctions (Fig. 5e). These results suggest that microtubule disruption destabilizes the actin cytoskeleton associated with cell-cell junctions and reduces transmission of rotational forces through AJs, even though it does not significantly alter ECad levels at cell-cell junctions. Taken together, we propose that microtubule disruption shifts the balance of force transmission by strengthening cell-substrate adhesion while weakening actin integrity at cell-cell junctions, thereby reversing collective rotation.

To elucidate how microtubule disruption affects the balance of force transmission between cell-cell and cell-substrate adhesions, we screened several microtubule-associated proteins whose depletion causes reversed collective rotation similar to that induced by nocodazole treatment and identified ACF7 as a candidate regulator (Fig. 6a,b; Extended Data Fig. 6a). ACF7, also known as MACF1, is an actin-microtubule cross-linking protein that coordinates the cytoskeletal networks and regulates both cell-cell and cell-substrate adhesions^14–16^. In particular, ACF7 has been shown to guide microtubule plus ends to FAs and to play important roles in microtubule dynamics and cellular adhesion^16^. ACF7 knockdown did not strongly affect nuclear rotation either in singly isolated cells or in colonies, indicating that ACF7 is dispensable for cell chirality (Fig. 6c; Extended Data Fig. 6b,c; Supplementary Videos 20 and 21). Nevertheless, nearly half of the colonies composed of ACF7-depleted cells exhibited reversed collective rotation, and the average collective rotation speed was approximately 2.5 degrees per hour counterclockwise direction (Fig. 6d; Supplementary Videos 22 and 23).

**Figure 6:**
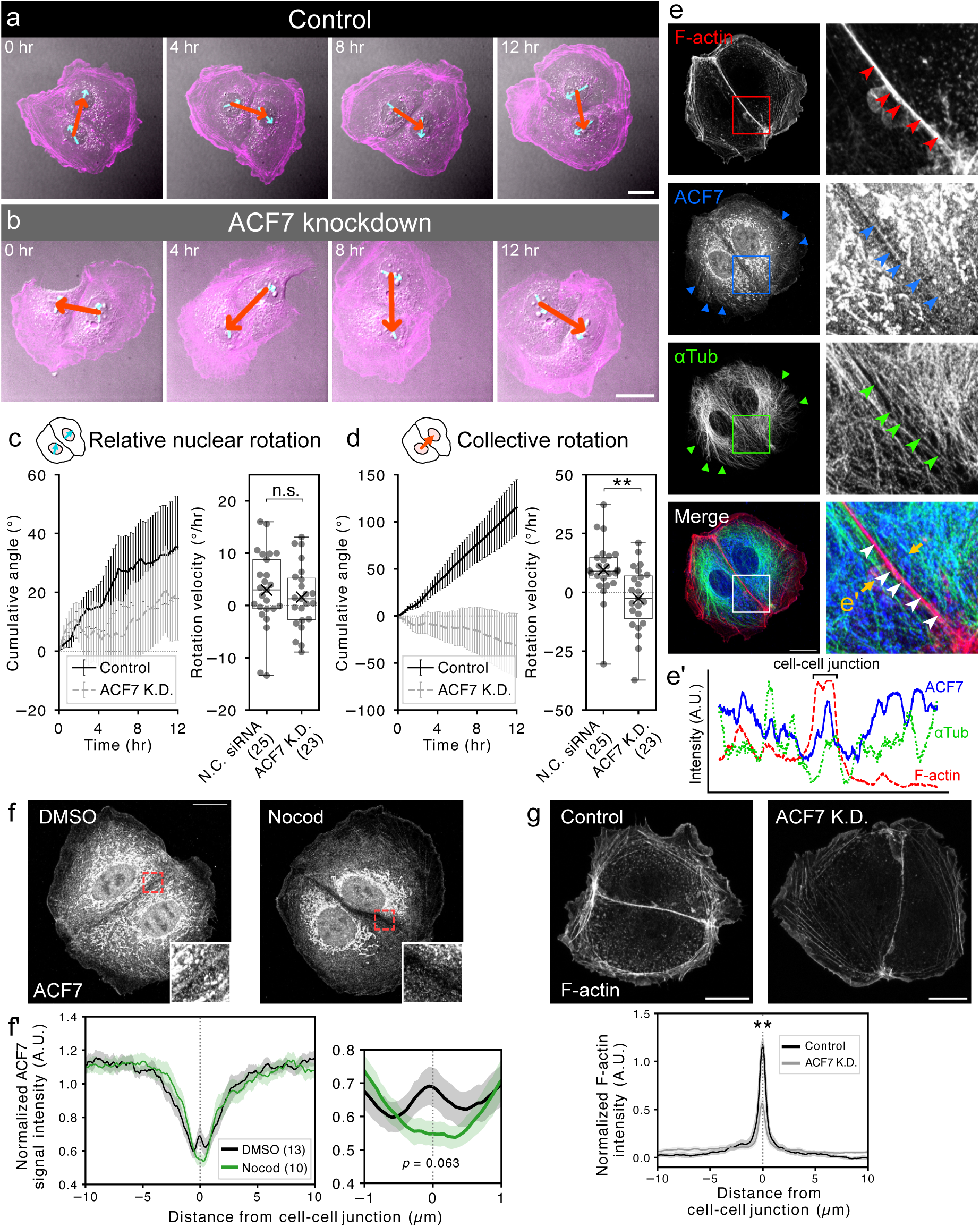
ACF7 knockdown reverses collective rotation by regulating F-actin integrity at cell-cell junctions. **a-b**, Representative time courses of paired Caco-2 cells treated with control (a) or ACF7 siRNA (b) are shown. Time-lapse images of DIC (gray) and Lifeact (magenta) are shown. Cyan and red arrows are the vectors used to measure nuclear rotation and collective rotation, respectively. **c-d**, Quantification of relative nuclear rotation (c) and collective rotation (d). Left: Time courses of cumulative rotation angles over 12 h. Black solid and gray dashed lines indicate cumulative rotation angles in control and ACF7 siRNA-treated cells, respectively. Error bars indicate standard errors. Right: Box plots of angular velocities calculated for 12 h of observation. Circles indicate angular velocities of individual cells. Crosses indicate average values. **e**, Representative image of F-actin (phalloidin), ACF7, and microtubule (anti-*α*tubulin) in a Caco-2 colony. Regions marked by boxes in the left panels are shown at higher magnification in the right panels. Blue and green arrowheads indicate peripheral ACF7 and microtubules, respectively. White arrowheads indicate cell-cell junctions where F-actin, ACF7, and microtubules colocalize. The immunofluorescence intensity profile shown in e’ was obtained by line scanning (width = 50 px) along the line connecting the tips of the yellow arrows in e. **f**, Loalization of ACF7 at cell-cell junctions. Top panels: Representative images of DMSO-treated cells (left) and nocodazole-treated cells (right). Insets show magnified views of the cell-cell junction regions indicated by red boxes. (f’) Left: Averaged profiles of normalized ACF7 intensity within 10 *μ*m of cell-cell junctions from multiple samples. Black and green lines indicate normalized signal intensities of DMSO-treated and nocodazole-treated cells, respectively. Right: Enlarged view of the region within 1 *μ*m of the junction. **g**, Localization of F-actin at cell-cell junctions. F-actin was visualized with phalloidin. Top: Representative images of cells treated with control (left) and ACF7 siRNA (right). Bottom: Averaged profiles of normalized F-actin intensity within 10 *μ*m of cell-cell junctions from multiple samples. Black, control; gray, ACF7 knockdown. Sample sizes are shown in parentheses. Scale bars: 20 *μ*m. n.s.: *p* > 0.05; **: *p* < 0.01 by *t*-test.

To investigate how ACF7 regulates cell-cell and cell-substrate force transmission, we next examined its subcellular localization. As previously reported^17^, ACF7 localized to microtubule tips at the cell periphery (Fig. 6e, blue and green triangles). Furthermore, we frequently detected colocalization of ACF7 and F-actin at cell-cell junctions (eight of 13 control colonies; Fig. 6e, red and blue arrowheads), although the ACF7 signal was relatively weak. Interestingly, at cell-cell junctions where ACF7 and F-actin colocalized, microtubule ends occasionally aligned with these sites (Fig. 6e, green arrowheads; Fig. 6e’). To test whether microtubules are required for junctional recruitment of ACF7, we inhibited microtubule polymerization with nocodazole and found that the ACF7 localization at the cell-cell junctions was frequently reduced or lost (eight of ten colonies; Fig. 6f,f’). These results suggest that microtubules recruit ACF7 to cell-cell junctions.

To determine whether ACF7 knockdown affects cell-cell force transmission in a manner similar to nocodazole treatment, we examined FAs and AJs in ACF7-depleted colonies. We first assessed FAs and found that ACF7-depleted colonies exhibited significantly more and larger FAs than control colonies (Extended Data Fig. 6d,e), consistent with a previous report^14^. Next, we examined AJs in ACF7-depleted colonies. As in nocodazole-treated colonies, ECad localization at cell-cell junctions did not change (Extended Data Fig. 6f,g). Meanwhile, reduced F-actin at cell-cell junctions, similar to that observed in nocodazole-treated colonies, was also detected in ACF7-depleted colonies (Fig. 6g), although F-actin detachment at cell-cell junctions was not frequently observed (one of ten colonies). These findings suggest that ACF7 helps maintain junctional F-actin through actin-microtubule coordination. Thus, ACF7 depletion may mimic microtubule disruption by enhancing FAs while weakening force transmission between cells, thereby leading to reversed collective rotation.

### Intracellular force transmission by actin-cadherin coupling

These observations prompted us to hypothesize that the association between F-actin and AJs acts as a mechanical clutch that modulates the transmission of cellular rotational force into intercellular mechanical coupling. To test this possibility, we used previously established *α*E-Catenin knockout (*α*ECat KO) Caco-2 cells^18^ and first examined their collective behavior. Strikingly, *α*ECat KO colonies exhibited reversed collective rotation (Fig. 7a; Supplementary Video 24). Approximately half of the colonies rotated counterclockwise, and because counterclockwise rotation was faster than clockwise rotation, the average collective rotation speed in *α*ECat KO colonies was about five degrees per hour in the counterclockwise direction (Fig. 7b). Despite this reversal of collective rotation, cell chirality remained intact. *α*ECat KO cells rotated clockwise under singly isolated conditions (Extended Data Fig. 7a,b; Supplementary Video 25), and relative nuclear rotation also remained clockwise in *α*ECat KO colonies (Fig. 7c), indicating that *α*ECat is dispensable for cell chirality. Consistent with the role of *α*ECat in physically linking ECad to the actin cytoskeleton^19^, F-actin was frequently detached from cell-cell junctions in *α*ECat KO colonies (Fig. 7d). These results suggest that actin anchoring at cell-cell junctions is crucial for transmitting cell-intrinsic rotational forces into collective rotation.

**Figure 7:**
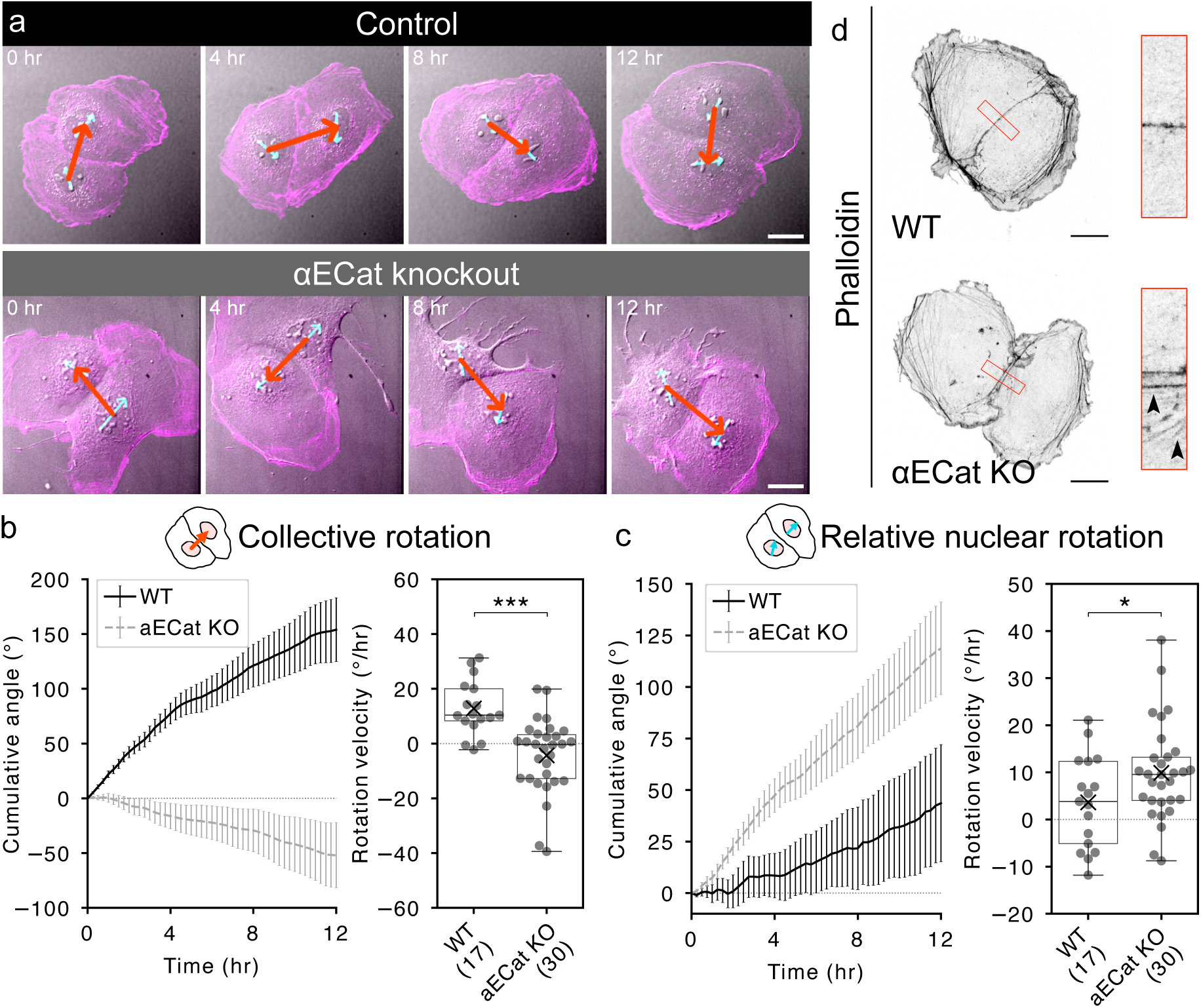
Disruption of actin-cadherin linkage reverses collective rotation. **a**, Representative time courses of wild-type (top) and *α*ECat KO Caco-2 cell pairs (bottom). Time-lapse images of DIC (gray) and Lifeact (magenta) are shown. Cyan and red arrows are the vectors used to measure nuclear rotation and collective rotation, respectively. **b-c**, Quantification of relative nuclear rotation (b) and collective rotation (c). Left panels: Time courses of cumulative rotation angles over 12 h. Black solid and gray dashed lines indicate cumulative rotation angles in wild-type and *α*ECat KO cells, respectively. Error bars indicate standard errors. Right panels: Box plots of angular velocities calculated for 12 h of observation. Circles indicate angular velocities of individual cells. Crosses indicate average values. Sample sizes are shown in parentheses. **d**, Localization of F-actin at cell-cell junctions visualized by phalloidin staining. Representative images of wild-type (top) and *α*ECat KO cells (bottom) are shown. Right panels show magnified views of the cell-cell junction regions indicated by red boxes. Arrowheads indicate detached F-actin. Scale bars: 20 *μ*m. *: *p* < 0.05; ***: *p* < 0.001 by *t*-test.

## Discussion

In this study, we show how homochiral cells collectively give rise to multicellular handedness. Using epithelial Caco-2 cells that exhibit clockwise cell chirality, we demonstrated that the left-right (LR) asymmetry of collective motion is determined by the balance of force transmission through cell-cell and cell-substrate adhesions. When chiral forces are transmitted preferentially through adherens junctions (AJs), cells gain traction at cell-cell interfaces, leading to collective rotation in the same direction as individual cell rotation. By contrast, when force transmission is biased toward focal adhesions (FAs), cells gain traction at the colony periphery, resulting in collective rotation opposite to single-cell rotation. Our theoretical model showed that this mechanical balance is sufficient to explain both the emergence and reversal of collective rotation without altering the intrinsic chirality of single-cells. This prediction was supported experimentally: inhibition of AJs with an E-Cadherin (ECad) inhibitory antibody induced reversal of collective rotation, whereas activation of FAs by FAK overexpression suppressed it. We further showed that microtubule disruption or ACF7 depletion also reversed collective rotation. Our results suggest that microtubules regulate intracellular force transmission by recruiting ACF7 to cell-cell junctions, where ACF7 maintains F-actin integrity and enables actomyosin-generated forces to be transmitted through AJs. Consistently, deletion of *α*E-catenin (*α*ECat), which anchors F-actin to ECad, also reversed collective rotation without affecting single-cell chirality. This finding further supports the idea that actin anchorage at AJs is essential for transmitting intracellular rotational forces into intercellular mechanical coupling. More broadly, our results suggest that the spatial arrangement of force-transmitting adhesions across distinct cellular interfaces is an important determinant of collective behavior. Such spatial organization allows forces arising from a common intracellular source to be routed through different external interfaces and thereby converted into distinct multicellular dynamics. Together, these findings show that mechanical transmission of actomyosin-generated forces, from intracellular actin dynamics to intercellular adhesions, provides a fundamental mechanism linking single-cell chirality to multicellular LR asymmetry across scales.

Our results further suggest that the balance between these spatially separated adhesive routes is actively regulated by the microtubule-ACF7 system. Consistent with previous *in vivo* and *in vitro* studies^20,21^, microtubule disruption impaired mechanical transmission to neighboring cells by weakening the connection between F-actin and AJs. Furthermore, supported by previous *in vitro* studies using truncated ACF7^22,23^, our results suggest that microtubules recruit ACF7 to the cell-cell boundary, where its actin-bundling activity helps maintain junctional actin integrity and thereby supports efficient force transmission through AJs. In parallel, ACF7 depletion increased focal adhesions, in agreement with previous findings that ACF7 guides microtubules along F-actin to focal adhesions and that its loss stabilizes FA-actin networks^14^. Together, these observations suggest that microtubules and ACF7 promote AJ-mediated force transmission while simultaneously preventing excessive stabilization of FAs. The weaker F-actin detachment phenotype at cell-cell junctions in ACF7-depleted colonies than in nocodazole-treated colonies may reflect adaptation to chronic depletion, in contrast to the acute disruption caused by nocodazole after colony formation. Direct analysis of ACF7 itself remains challenging because of its large size (> 600 kDa) and multiple splice variants, and was therefore beyond the scope of this study. Our study highlights microtubule-dependent regulation of adhesion balance as an important mechanism that determines where intracellular forces are anchored during collective cell behavior.

Importantly, our study provides an alternative framework in which multicellular LR-asymmetry is not necessarily a direct reflection of cell chirality. Several groups, including ours, have reported reversals of multicellular LR asymmetry *in vivo* and *in vitro*^5,24–30^, which have often been attributed to reversal of cell chirality^5,24,26–28,30,31^. However, this interpretation leaves unresolved the fundamental question of how cell chirality itself would reverse. Instead, our mechanical framework suggests that multicellular LR asymmetry is determined not by cell chirality alone, but by how chiral forces are transmitted through cell-cell and cell-substrate adhesions. In this view, reversal and randomization of multicellular LR asymmetry can arise from changes in the balance of force transmission without requiring inversion of single-cell chirality. Previous reports are consistent with this framework: reduced cell-cell adhesion is associated with a higher frequency of LR inversion in multicellular endothelial systems^32^ and in MDCK cells^33^, and enlargement of FAs also reverses multicellular LR asymmetry^34^. Furthermore, *in vivo* studies have shown that the deletion or depletion of DECad randomizes the LR asymmetric rotation of the gut and male genitalia in *Drosophila*^5,28,35^. Overall, these observations suggest that the balance of force transmission through cell-cell and cell-substrate adhesions is a general mechanism underlying multicellular LR asymmetry across biological systems.

Badih *et al.* and Im *et al.* also investigated clockwise-biased two-cell collective rotation using a confined hUVEC system and suggested that contractile force strength determines collective rotation direction^36,37^. Although our cells also exhibited clockwise collective rotation, our system differs from the hUVEC system in the epithelial origin of Caco-2 cells, the presence of intrinsic single-cell rotation, and the absence of substrate confinement. Furthermore, in the hUVEC model, blebbistatin increases the frequency of clockwise collective rotation, whereas in our system, blebbistatin leads to randomized rotation. These differences suggest that multiple distinct mechanisms can generate collective rotation. Future work integrating our molecular findings with quantitative force measurements and theoretical modeling will be important for establishing a more complete understanding of how the balance of force transmission between cell-cell and cell-substrate adhesions shapes collective cell dynamics.

Our work focused on two-cell colonies, and further studies are needed to fully understand the relationship between cell chirality and LR asymmetry in confluent cell sheets. Because two-cell colonies lack tricellular junctions and stable tight junctions, it remains unclear how the absence of these structures affects chiral collective rotation driven by cellular rotational forces. Additionally, recent work has shown that hUVECs cultured as confluent tubular structures can exhibit twisting behavior^30^. Whether the mechanisms underlying this tube twisting are similar to those governing collective rotation in two-cell colonies remains an open question and warrants further investigation.

## Methods

### Cell culture, live imaging, and drug treatment

Caco-2 cells obtained from ATCC were cultured in DMEM/Ham’s F-12 (FUJIFILM Wako Pure Chemical Corp. #048-29785) supplemented with 10% fetal bovine serum (FBS), 1x GlutaMAX Supplement (ThermoFisher #35050061), and 1% penicillin and streptomycin (Nacalai Tesque #26253-84) at 37 °C in 5% CO_2_. For live imaging of actin, we used previously established Lifeact-RFP-expressing Caco-2 cells^10^, and observed them on a 35 mm collagen type I-coated dishes (IWAKI #4970-011). *α*ECat KO Caco-2 cells expressing Lifeact-RFP were previously established^18^. Live imaging was performed as previously described^10^.

Drugs were added to the medium 2-4 h before observation for single-cell rotation assays and one day before observation for collective rotation assays. Because cell division transiently interrupts both single-cell and collective rotations and affects quantification, 0.2 - 0.6 µM of aphidicolin (FUJIFILM, #011-09811) was also added to arrest cell cycles before live imaging of single-cells and two-cell colonies. The following reagents were used: 3 µM blebbistatin (SIGMA, #B0560-1MG); 50 µM nocodazole (SIGMA, #M1404); 1 µg/mL SHE78-7 (TAKARA, #M126); 1 µg/mL mouse normal IgG (SantaCruz, #sc-2025); 0.2% DMSO (negative control).

### Transfection

For gene knockdown, 2.67 × 10) cells were seeded in a 60 mm dish on day 0, and 8.5 µM siRNA was transfected on day 1 using Lipofectamine RNAiMAX (Invitrogen #13778030). Live imaging was performed on day 5. The following siRNAs were used: ACF7 (Invitrogen, #HSS53618496); Medium GC negative control (Invitrogen #12935300).

To construct FAK::GFP and GFP overexpression vector, the LifeAct-mEmerald region of the previously described pLVSIN-EF1a-Lifeact-mEmerald-IRES-pur plasmid^10^ was replaced using In-Fusion cloning (Takara). Plasmids were introduced into cells using Lipofectamine 3000 Reagent (Invitrogen, #L3000001), according to the manufacturer’s protocol.

### Immunofluorescence staining and imaging

Cells were fixed and permeabilized at 37°C with 4% of paraformaldehyde-PBS and 0.2% Triton-X-100, and then washed three times with 1 × PBS. Fixed cells were blocked with Blocking One (Nacalai tesque, #03953-95) and incubated for 1 h at room temperature or overnight at 4°C with primary antibodies: rabbit anti-myosin II (Sigma-Aldrich, #M8064, dilution 1:500); mouse anti-phosphorylated myosin light chain 2 (Ser19) (Cell Signaling, #3675S, dilution 1:200); rabbit anti-pFAK (Thermo Fisher, #44-624G, dilution 1:250); rat anti-*α*-tubulin (Santa Cruz, #sc-53029, dilution 1:500); mouse anti-E-Cadherin (BD biosciences, #610181, dilution 1:100); rabbit anti-ACF7 (a generous gift of Dr. Wenxiang Meng^17^, dilution 1:1000).

Cells were then incubated for 2 h at room temperature with appropriate AlexaFluor-conjugated secondary antibodies (Molecular Probes, dilution 1:400). Actin was stained with phalloidin (Invitrogen, #A12379 for Alexa Fluor 488 phalloidin or #A12380 for Alexa Fluor 568 phalloidin). CanGetSignal Solution B (TOYOBO, #NKB-601) was used to dilute antibodies.

Fluorescence images were acquired using a laser scanning confocal microscope (Zeiss, LSM880). Images were processed with Fiji.

### Analysis of cell rotation

We quantified nuclear rotation in the laboratory frame, *θ_lab_*, by manually tracking two nuclear speckles in each cell on DIC images using Manual Tracking plugin in Fiji. The nuclear rotation angle was defined as the angle of the line connecting two nuclear speckles, and rotational dynamics were analyzed using Python3. To calculate nuclear rotation during collective rotation, we used the mean of the two nuclear rotation angles in each colony. To quantify the angle of collective rotation, *Θ*, we measured the angle of the line connecting the centers of the two nuclei. The midpoint of the line connecting the two nuclear speckles was used to define the nuclear center. Nuclear rotation in the colony frame, *θ_cell_*, was calculated by subtracting *Θ* from *θ_lab_*. For single-cell rotation, nuclear rotation in the laboratory frame and in the colony frame were treated as identical. Sixteen-hour live-imaging movies were analyzed, and data from the first 12 h were used to calculate the angular velocity of collective rotation *Ω* and nuclear rotation *ω_lab_*. In this paper, *θ_lab_* and *ω_lab_* are presented as *θ* and *ω*, respectively. Cells that underwent cell division or cell death, cells whose nuclei moved out of the field of view, and cells that adhered to cells migrating into the field of view from outside the image were excluded from the analysis.

For analysis of larger colonies (Extended Data Fig. 2), nuclear center positions were manually annotated. Cell positions and cell division events were then automatically measured using TrackMate plugin in Fiji. To minimize the influence of mitosis on the measurement of colony rotation, the division frame and all frames within ±30 min of each cell division event were excluded from the analysis. Angular velocity around the colony center of mass was then calculated in Python 3 only for colonies that exhibited continuous rotation for at least 3.5 h without cell division during the analyzed period.

Statistical analyses were performed using scikit-posthocs (https://github.com/maximtrp/scikit-posthocs).

### Numerical simulation of the theoretical model

We modeled the rotating cells using polygons. The equation of motion for the vertices of the polygons is given by

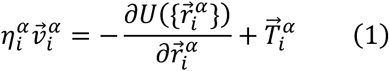

where the left-hand side is the frictional force reflecting the cell-substrate adhesion with coefficient 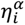 and velocity 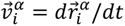 of vertex *i* of cell *α* with the positional vector 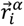. The first term on the right-hand side is the potential force with potential function 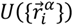, and the second term is the active force reflecting the cell chirality with 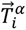 working along the periphery of a cell. The potential function 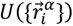 consists of area elasticity, line elasticity, bending rigidity, and cell-cell adhesion. For further details of our theoretical model, see Supplementary Notes.

Among the angular velocity 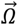 of the colony, the cell-cell adhesion force 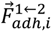 exerted on vertex *i* of cell 1 by cell 2, and the velocity of vertex *i*, 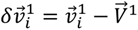, relative to the centroid velocity of cell 1, 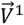, the following relation holds:

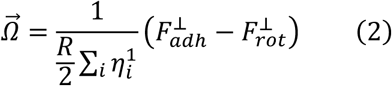

with

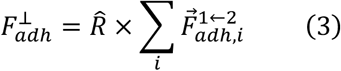

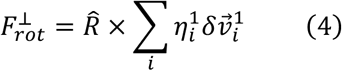

where *R* and *R̂* are the distance and the unit vector between the two cell centroids, respectively. 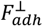 is the contribution from the cell-cell adhesion force exerted on cell 1 by cell 2, and 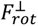 is the cell-substrate frictional force of cell 1 contributed from the rotation of cell 1. The balance between these two forces determines the direction (sign) of the angular velocity 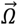. We also notice that the only tangential components of these forces (i.e., the components perpendicular to the line connecting the two cell centroids) contribute to the collective rotation.

A boundary of clockwise and counterclockwise phases 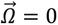 and contour lines for collective rotation velocity 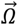 are estimated with the cubic interpolation function implemented in scipy library. The estimated phase boundary was fitted with the orthogonal distance regression function implemented in scipy library.

### Image analysis of boundary signals and focal adhesions

The cell-cell boundary was manually annotated, and boundary images of ECad, phalloidin, and ACF7 were computationally straightened using “Straighten” function in Fiji. A square region was cropped from the center of each straightened boundary image, and signal intensity was averaged along the cell-cell boundary. The signal was normalized by subtracting the mean intensity of the five micrometer regions at both edges of the cropped image. The peak value of the averaged signal intensity was then quantified.

To measure the number and area of focal adhesions, pFAK images were binarized using a custom ImageJ macro (https://gist.github.com/ishibaki/603d11edc6b535699f987065b5f3b3da). The number and area of FAs were then quantified using “Analyze Particles” function in Fiji. Statistical analyses were performed using scikit-posthocs.

### Immunoblotting

Cells were lysed in RIPA buffer on day 5 and proteins were extracted. Proteins were separated by SDS-PAGE on 5% polyacrylamide gels and transferred to 0.45 µm PVDF membranes (FUJIFILM, #038-25661) at 300 mA for 2.5 h. Membranes were then blocked for 20 min with Blocking One and incubated at room temperature with the following primary antibodies: mouse anti-FAK (Santa Cruz, #sc-271126, dilution 1:1000); rabbit anti-pFAK (Thermo Fisher, #44-624G, dilution 1:1000); mouse anti-ACF7 (Abnov, #H00023499-A01, dilution 1:500); mouse anti-GAPDH (Santa Cruz, #sc-166574, dilution 1:1000). After primary antibody incubation, membranes were processed for ECL Select detection using appropriate HRP-conjugated second antibodies: HRP anti-mouse IgG (Invitrogen, #T20912, dilution 1:5000); HRP anti-rabbit IgG (Invitrogen, #T20926, dilution 1:5000). Primary and secondary antibodies were diluted in CanGetSignal Solutions 1 and 2 (TOYOBO, #NKB-101T), respectively.

## Acknowledgments

We thank Masatoshi Takeichi for generously providing the Caco-2 cells used in this study. We thank Wenxiang Meng, Yuko Kiyosue-Mimori, and Yasushi Okada for sharing antibodies and reagents, and Kyogo Kawaguchi and RIKEN KBiF for providing equipment. We are grateful to Yu-Chiun Wang, Fumio Motegi, and Yukako Nishimura for critical reading of the manuscript, and to Hiroyuki Iida, and Kentaro Yamamoto for valuable comments on the manuscript. We thank Masayuki Hayakawa, Biplab Bhattacherjee, Vivek Semwal, Motohiro Fujiwara, Debjyoti Majumdar, Abhinandan Angra, Takanobu A. Kato, Yohsuke T. Fukai, Kyosuke Adachi, and colleagues at RIKEN BDR for fruitful discussions. This research was supported by Japan Society for the Promotion of Science (JSPS) KAKENHI (23K14186, 22KJ3145, and 22J01009 to TI; 23K16999 to GO; 22H05170, 23H02455, 25K22443, and 26H02374 to TS), Japan Science and Technology Agency (JST) ACT-X (JPMJAX2423 to TI), and JST CREST (JPMJCR1852 to TS). TI, RN, and GO were supported by RIKEN Program for Young Scientists.

## Author contributions

T.I.: Conceptualization, Data curation, Formal analysis, Funding acquisition, Investigation, Methodology, Project administration, Resources, Software, Supervision, Validation, Visualization, Writing – Original Draft, Writing – Review & Editing.

R.N.: Data curation, Formal analysis, Investigation, Methodology, Software, Visualization, Writing – Review & Editing.

G.O.: Data curation, Formal analysis, Funding acquisition, Writing – Review & Editing.

N.T.: Data curation, Formal analysis, Investigation.

T.S.: Conceptualization, Funding acquisition, Project administration, Supervision, Writing – Review & Editing.

