## Extended Data Figures for "Force transmission balance through adhesions determines multicellular handedness"

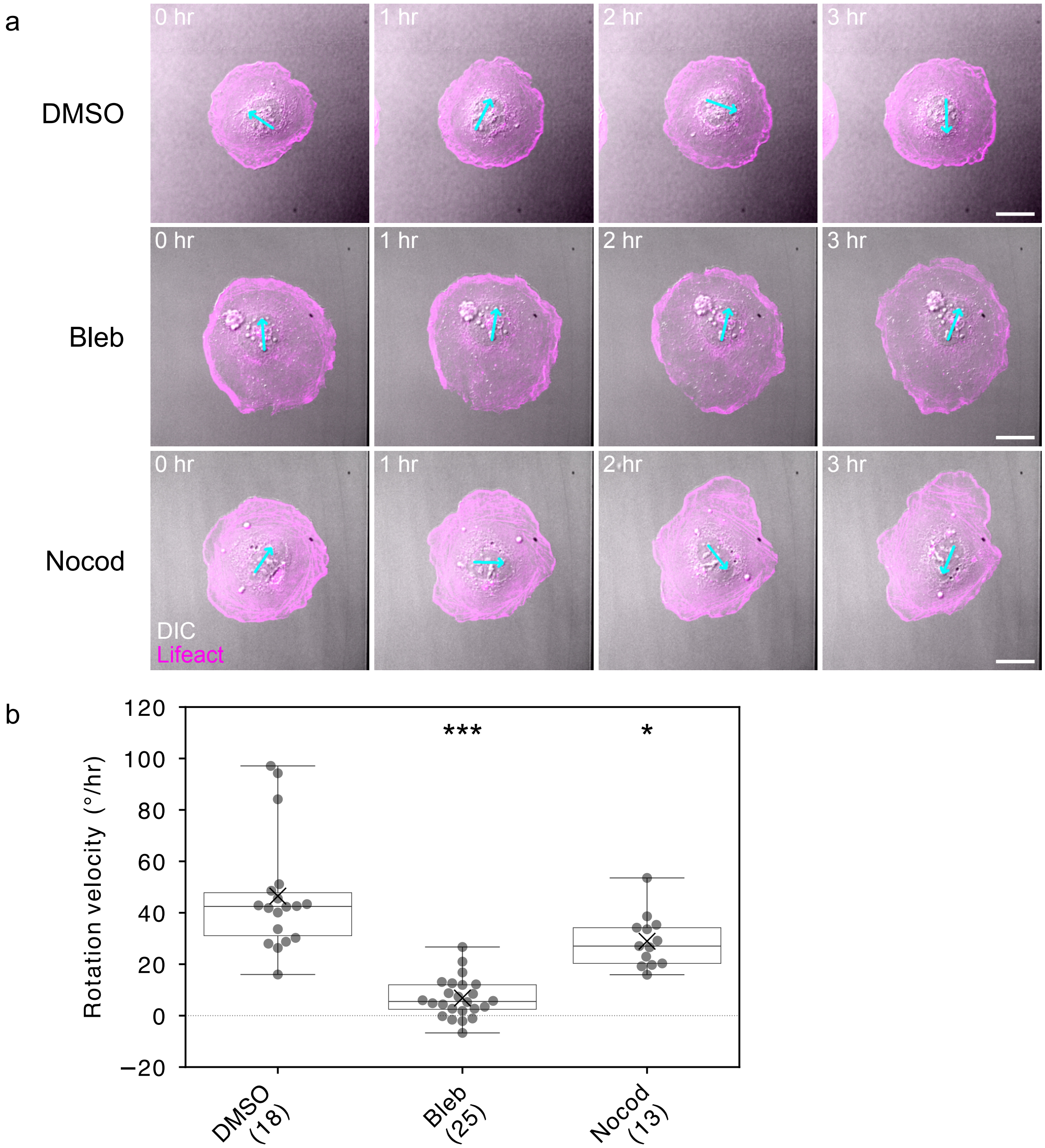


**Extended Data Figure 1:** Nuclear rotation of singly isolated cells. **a**, Representative time courses of singly isolated Caco-2 cells treated with DMSO (top), blebbistatin (middle), and nocodazole (bottom). Time-lapse images of DIC (gray) and Lifeact-RFP (magenta) are shown. Cyan arrows are the vectors connecting two nuclear speckles used to measure nuclear rotation. Scale bars: 20 $\mu$m. **b**, Box plot of angular velocity of clockwise nuclear rotation for 12 h. Circles and crosses indicate individual values and average values, respectively. Sample sizes are shown in parentheses. *: $p < 0.05$; ***: $p < 0.001$ by $t$-test followed by multiple pairwise comparisons with Bonferroni's correction.

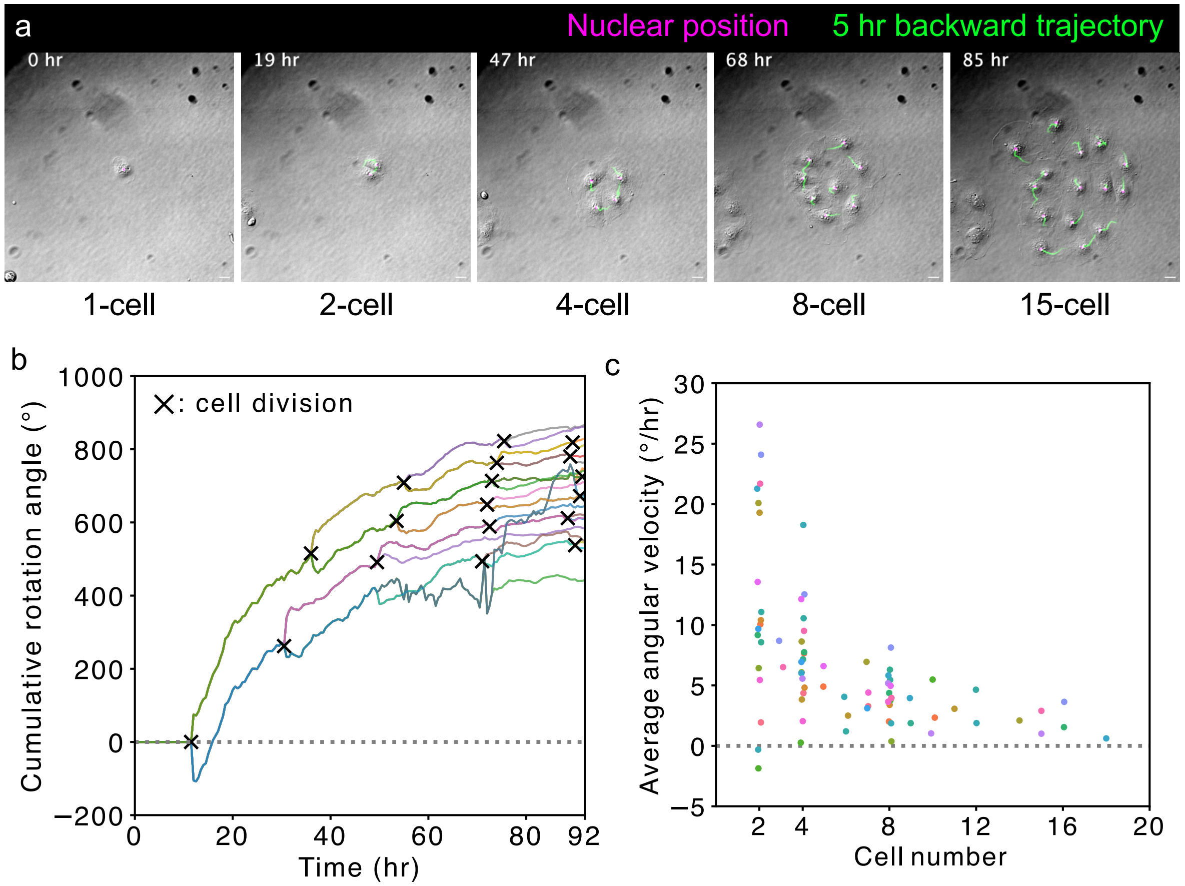


**Extended Data Figure 2:** Collective rotation of Caco-2 colonies containing two or more cells. **a**, Representative time courses of a Caco-2 colony showing clockwise collective rotation. Time-lapse DIC images are shown. Magenta circles indicate nuclear positions, and green lines indicate trajectories over the preceding 5 h. Scale bar: 20 $\mu$m. **b**, Cumulative angle of clockwise collective rotation for the colony shown in (a). Crosses indicate time points of cell division. Different colors indicate the time courses of individual cells within the colony. **c**, Dependence of collective rotation angular velocity on colony size. Colonies derived from the same mother cell are shown in the same color.

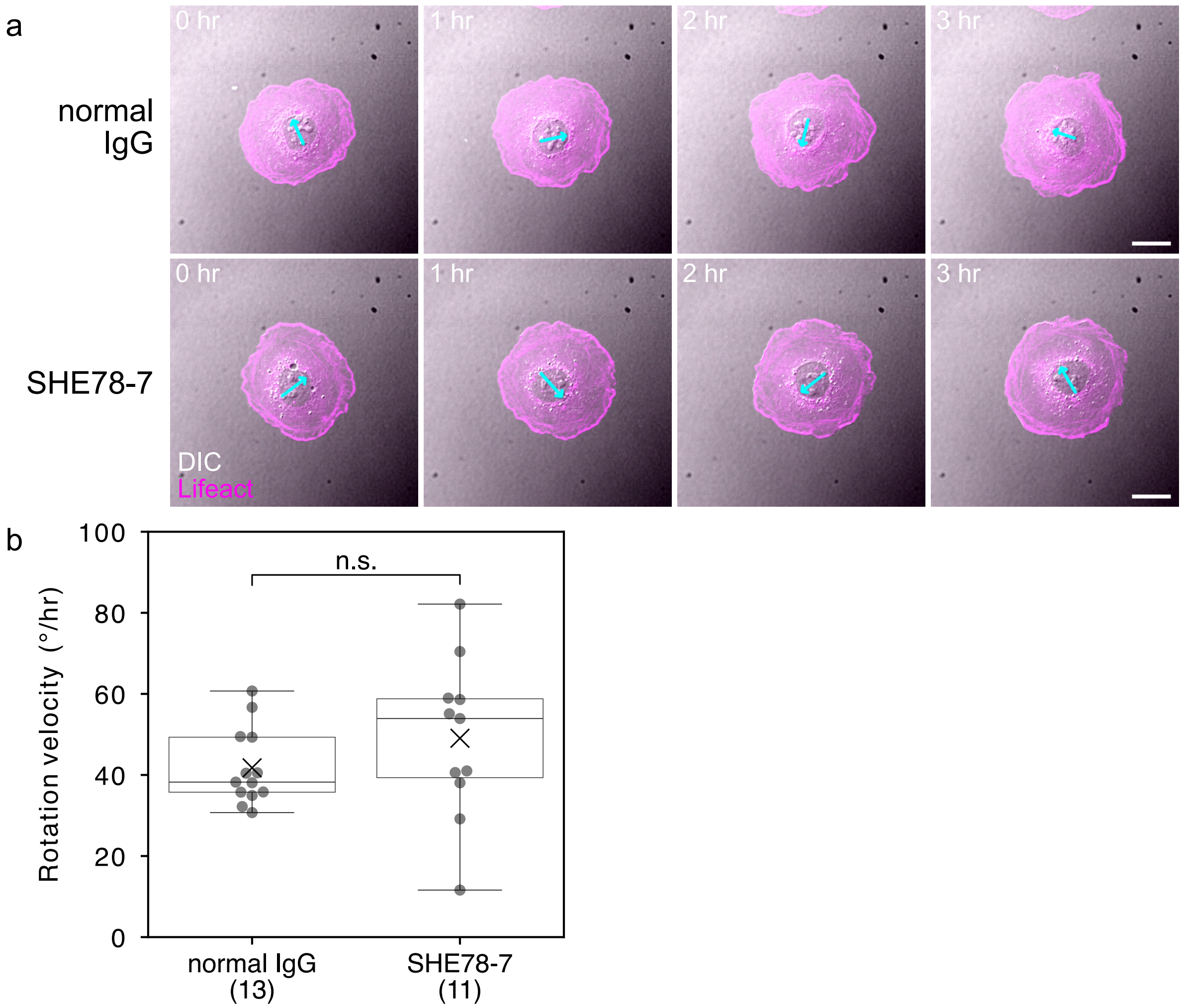


**Extended Data Figure 3:** Nuclear rotation of singly isolated cells under inhibitory antibody treatment. **a**, Representative time courses of singly isolated Caco-2 cells treated with normal mouse IgG (top) or SHE78-7, an inhibitory antibody against E-cadherin (bottom). Time-lapse images of DIC (gray) and Lifeact-RFP (magenta) are shown. Cyan arrows are the vector connecting two nuclear speckles used to measure nuclear rotation. Scale bars: 20 $\mu$m. **b**, Box plot of angular velocity of nuclear rotation for 12 h of observation. Circles and crosses indicate individual values and average values, respectively. Sample sizes are shown in parentheses. n.s.: $p > 0.05$ by $t$-test.

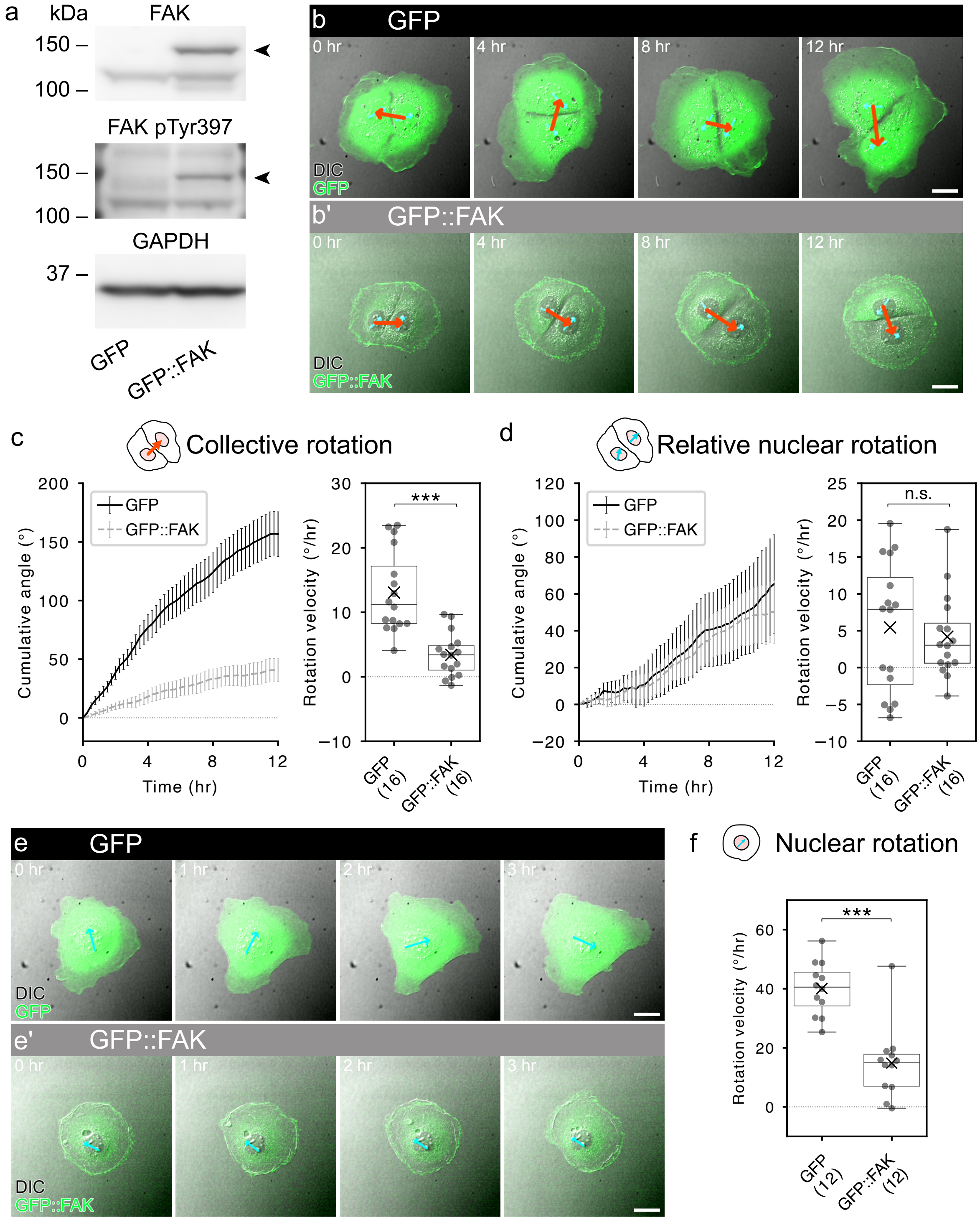


**Extended Data Figure 4:** FAK overexpression inhibits single-cell and collective rotation. **a**, Immunoblot detection of FAK, pFAK (pTyr397), and GAPDH in Caco-2 cells. **b**, Representative time courses of paired Caco-2 cells overexpressing GFP (b) and GFP::FAK (b'). Time-lapse images of DIC (gray) and GFP or GFP::FAK (green) are shown. Cyan and red arrows are the vectors used to measure nuclear rotation and collective rotation, respectively. **c-d**, Quantification of collective rotation (c) and relative nuclear rotation (d). Left panels: Time courses of cumulative rotation angles over 12 h. Black and gray lines indicate cumulative rotation angles in GFP- and GFP::FAK-overexpressing cells, respectively. Error bars indicate standard errors. Right panels: Box plots of angular velocities calculated for 12 h of observation. Circles indicate angular velocities of individual cells. Crosses indicate average values. **e**, Representative time courses of singly isolated Caco-2 cells overexpressing GFP (e) and GFP::FAK (e'). Time-lapse images of DIC (gray) and GFP or GFP::FAK (green) are shown. Cyan arrows are the vectors used to measure nuclear rotation. **f**, Quantification of nuclear rotation in singly isolated Caco-2 cells. Box plot of angular velocity calculated for 12 h of observation. Circles indicate angular velocities of individual cells. Crosses indicate average values. Scale bars: 20 $\mu$m. n.s.: $p > 0.05$; ***$p < 0.001$ by *t* test.

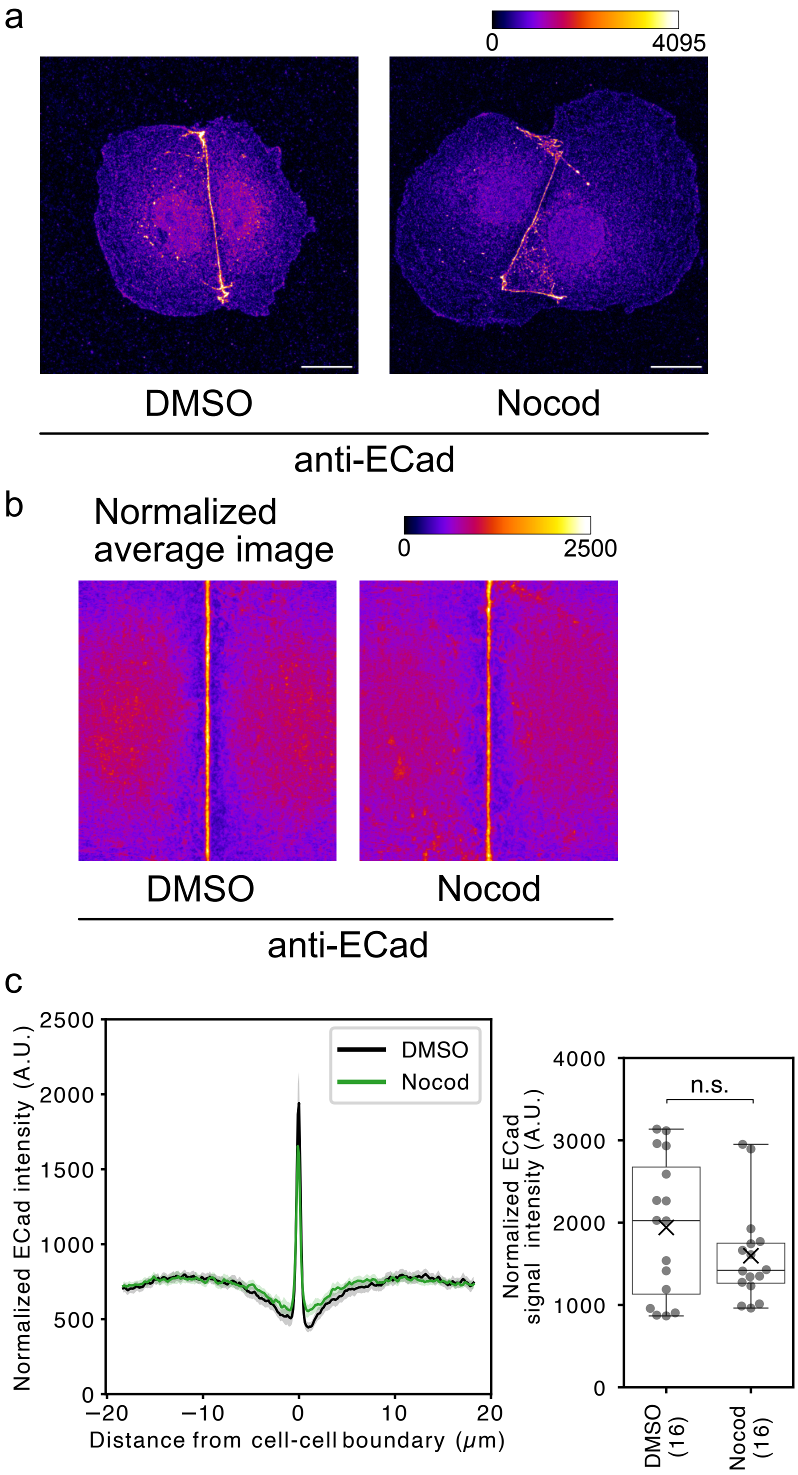


**Extended Data Figure 5:** E-cadherin levels at cell-cell junctions. **a**, Representative images of anti-E-cadherin staining in DMSO-treated (left) and nocodazole-treated cells (right). Scale bars: 20 $\mu$m. **b**, Average heatmap images of junctional E-cadherin. **c**, Quantification of normalized E-cadherin intensity. Left: Line plot of normalized average intensity of E-cadherin. Right: Box plot of normalized average intensity of E-cadherin at cell-cell junctions. Circles and crosses indicate individual values and average values, respectively. Sample sizes are shown in parentheses. n.s.: $p > 0.05$ by $t$-test.

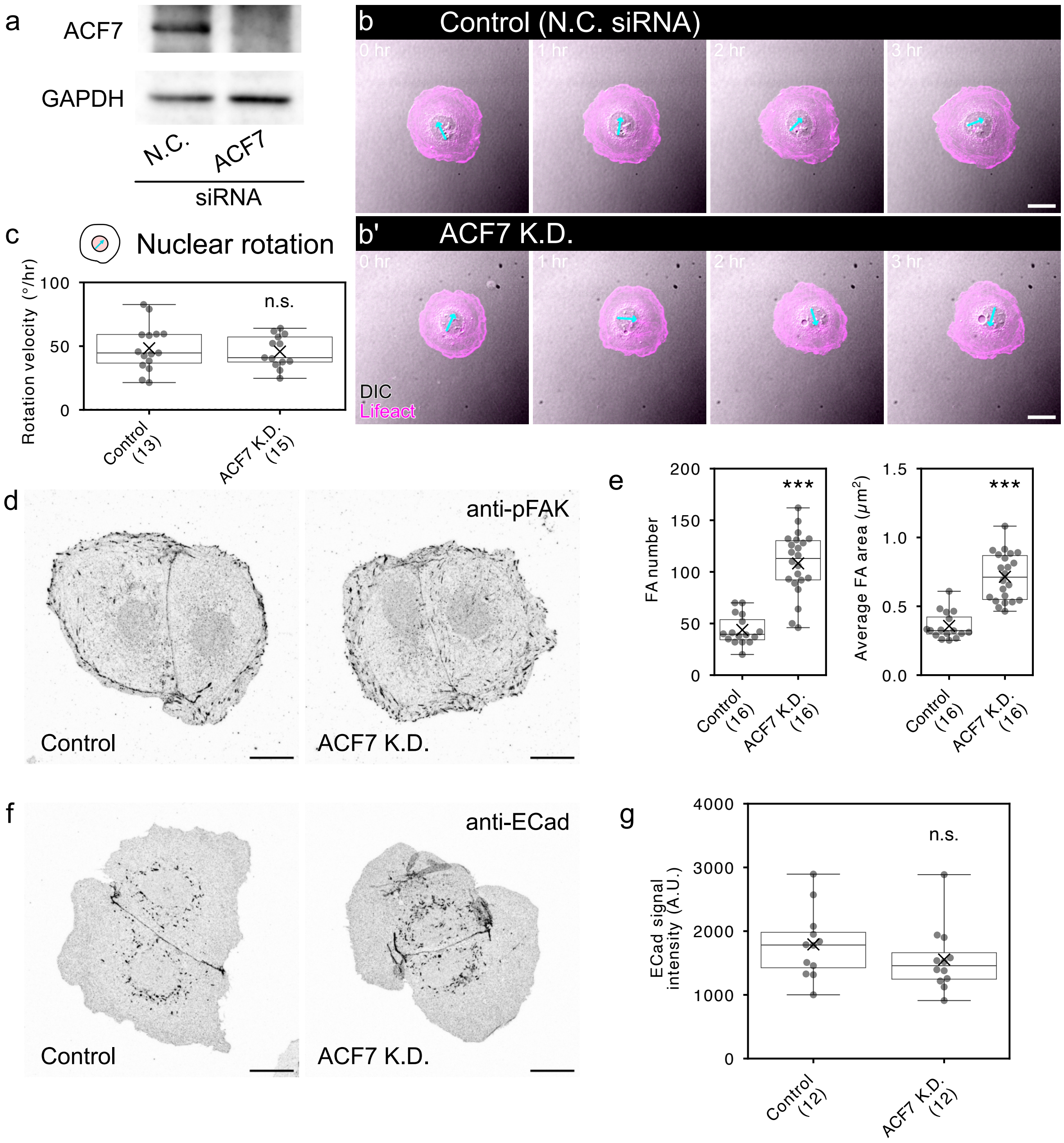


**Extended Data Figure 6:** Cell chirality and cell adhesions in ACF7-depleted cells. **a**, Immunoblot detection of ACF7 and GAPDH in Caco-2 cells. **b**, Representative time courses of singly isolated control (b) and ACF7-depleted cells (b'). Time-lapse images of DIC (gray) and Lifeact-RFP (magenta) are shown. Cyan arrows are the vector connecting two nuclear speckles used to measure nuclear rotation. **c**, Box plot of angular velocity of nuclear rotation for 12 h of observation. Circles and crosses indicate individual values and average values, respectively. n.s.: $p > 0.05$ by *t*-test. **d**, Representative anti-pFAK staining images of control (left) and ACF7-depleted cells (right). **e**, Box plots of the number of pFAK speckles (left) and their average area (right). ***: $p < 0.001$ by Mann-Whitney's *U*

test. **f**, Representative anti-E-cadherin staining images of control (left) and ACF7-depleted cells (right) are shown. **g**, Box plot of E-cadherin intensity at cell-cell junctions. n.s.: $p > 0.05$ by $t$-test. Sample sizes are shown in parentheses. Scale bars: 20 $\mu$m.

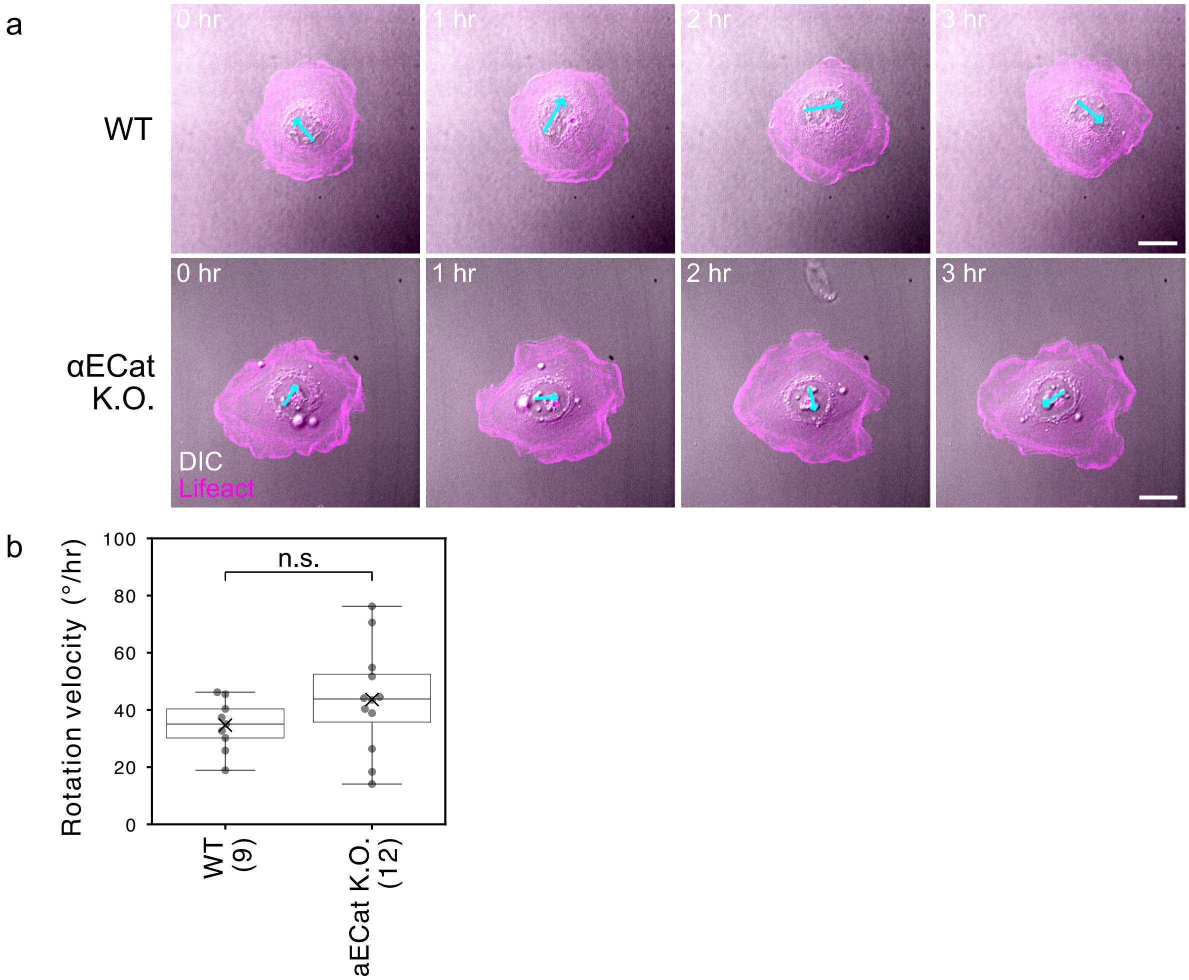


**Extended Data Figure 7:** Nuclear rotation of singly isolated $\alpha$ECat knockout cells. **a**, Representative time courses of singly isolated wild-type (top) and $\alpha$ECat knockout cells (bottom). Time-lapse images of DIC (gray) and Lifeact-RFP (magenta) are shown. Cyan arrows are the vector connecting two nuclear speckles used to measure nuclear rotation. Scale bars: 20 $\mu$m. **b**, Box plot of angular velocity of nuclear rotation for 12 h of observation. Circles and crosses indicate individual values and average values, respectively. Sample sizes are shown in parentheses. n.s.: $p > 0.05$ by $t$-test.
