## Supplementary Materials for "Force transmission balance through adhesions determines multicellular handedness"

### Supplementary Material

**Supplementary Figure**

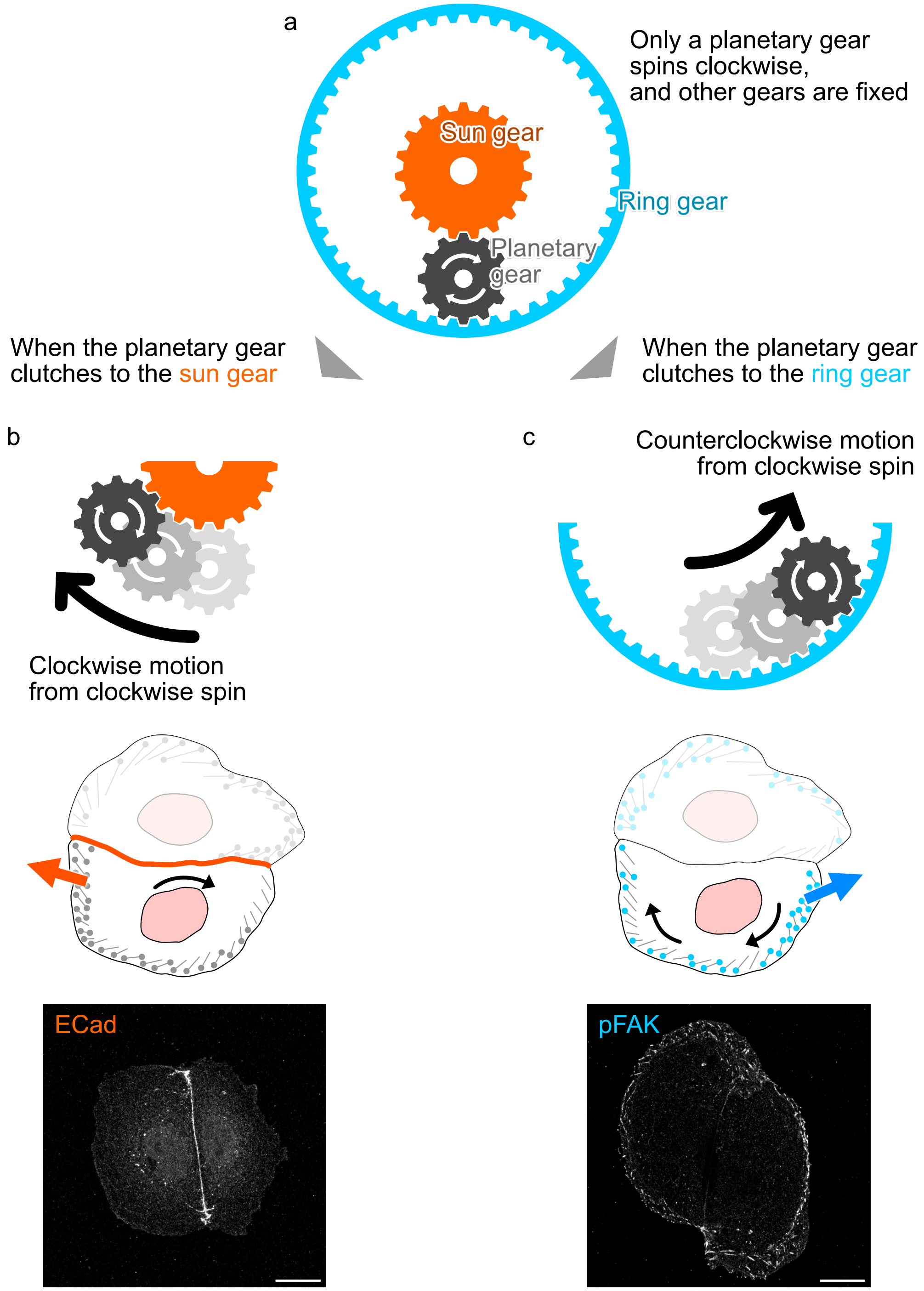

**Supplementary Figure 1:** Conceptual analogy between a planetary gear train and bidirectional collective rotation. **a**, Schematic illustration of a simplified planetary gear train. A planetary gear train typically consists of a central sun gear (orange), one or more planetary gears (black) that revolve around the sun gear, and an outer ring gear (blue) with internal teeth. The output direction depends on which gear is fixed or coupled; thus, a planetary gear train can generate either forward or reverse motion from the same input rotation. **b**, When the planetary gear is coupled to the sun gear, clockwise spin results in clockwise orbital motion of the planetary gear (top), analogous to clockwise collective rotation of a two-cell Caco-2 colony mediated by cell-cell adhesion (middle). Bottom, representative confocal image of E-cadherin (ECad) immunostaining, illustrating cell-cell adhesion. **c**, When the planetary gear is coupled to the outer ring gear, the same clockwise spin produces counterclockwise orbital motion of the planetary gear (top), analogous to counterclockwise collective rotation of a two-cell Caco-2 colony mediated by cell-substrate adhesion (middle). Bottom, representative confocal image of phosphorylated focal adhesion kinase (pFAK) immunostaining, illustrating cell-substrate adhesion. Scale bars: 20 $\mu$m.

### Supplementary Videos

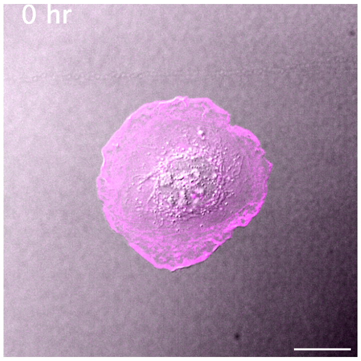

**Supplementary Video 1:** Clockwise rotation of a single Caco-2 cell. DIC and Lifeact-RFP signals are shown in gray and magenta, respectively. Images were acquired every 15 minutes for 12 h. Scale bar, 20 $\mu$m. Display rate, 20 frames/sec.

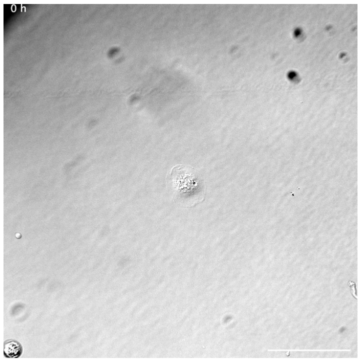

**Supplementary Video 2:** Growth of a multicellular Caco-2 colony. DIC images are shown. Images were acquired every 30 minutes for 92 h. Scale bar, 200 $\mu$m. Display rate, 20 frames/sec.

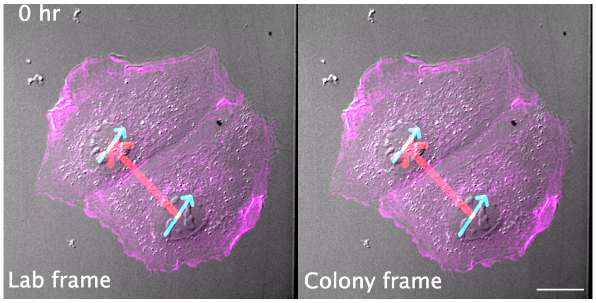

**Supplementary Video 3:** Clockwise collective rotation of a two-cell Caco-2 colony. DIC and Lifeact-RFP signals are shown in gray and magenta, respectively. The left and right panels show motion in the laboratory frame and colony frame, respectively. Cyan and red arrows indicate vectors used to measure nuclear rotation and collective rotation. Images were acquired every 15 minutes for 12 h. Scale bar, 20 $\mu$m. Display rate, 20 frames/sec.

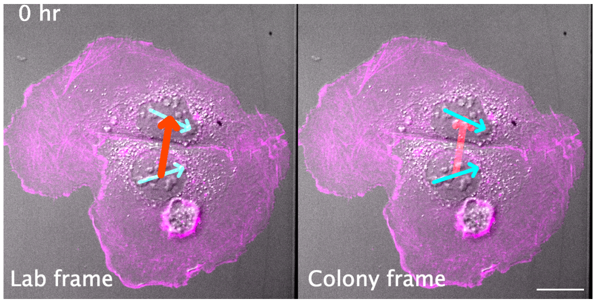

**Supplementary Video 4:** Arrested collective rotation of a blebbistatin-treated two-cell Caco-2 colony. DIC and Lifeact-RFP signals are shown in gray and magenta, respectively. The left and right panels show motion in the laboratory frame and colony frame, respectively. Cyan and red arrows indicate vectors used to measure nuclear rotation and collective rotation. Images were acquired every 15 minutes for 12 h. Scale bar, 20 $\mu$m. Display rate, 20 frames/sec.

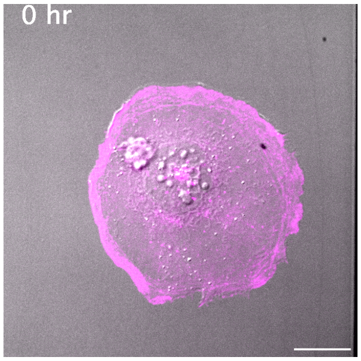

**Supplementary Video 5:** Arrested nuclear rotation of a blebbistatin-treated single Caco-2 cell. DIC and Lifeact-RFP signals are shown in gray and magenta, respectively. Images were acquired every 15 minutes for 12 h. Scale bar, 20 $\mu$m. Display rate, 20 frames/sec.

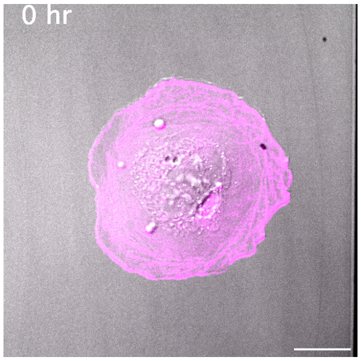

**Supplementary Video 6:** Clockwise rotation of a nocodazole-treated single Caco-2 cell. DIC and Lifeact-RFP signals are shown in gray and magenta, respectively. Images were acquired every 15 minutes for 12 h. Scale bar, 20 $\mu$m. Display rate, 20 frames/sec.

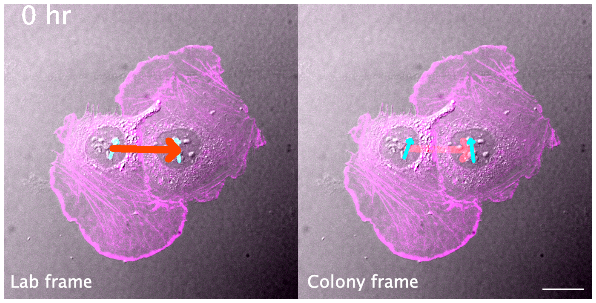

**Supplementary Video 7:** Counterclockwise collective rotation of a nocodazole-treated two-cell Caco-2 colony. DIC and Lifeact-RFP signals are shown in gray and magenta, respectively. The left and right panels show motion in the laboratory frame and colony frame, respectively. Cyan and red arrows indicate vectors used to measure nuclear rotation and collective rotation. Images were acquired every 15 minutes for 12 h. Scale bar, 20 $\mu$m. Display rate, 20 frames/sec.

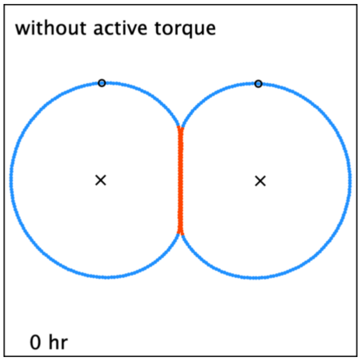

**Supplementary Video 8:** Clockwise collective rotation of a simulated two-cell colony under the control condition. Orange and blue circles indicate vertices associated with cell-cell adhesion and cell-substrate adhesion, respectively. Large black circles indicate vertex 0. Crosses indicate cell centers. The following parameters were used: $K_t = 0.035$; $K_{adh} = 7.0 \times 10^{-7}$; $\eta = 1.5$.

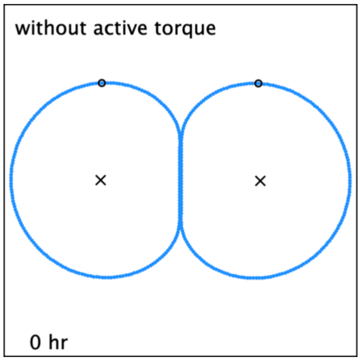

**Supplementary Video 9:** Counterclockwise collective rotation of a simulated two-cell colony with reduced cell-cell adhesion. Orange and blue circles indicate vertices associated with cell-cell adhesion and cell-substrate adhesion, respectively. Large black circles indicate vertex 0. Crosses indicate cell centers. The following parameters were used: $K_t = 0.035$; $K_{adh} = 3.4 \times 10^{-7}$; $\eta = 1.5$.

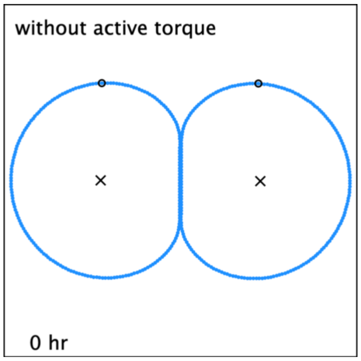

**Supplementary Video 10:** Slower clockwise collective rotation of a simulated two-cell colony with increased cell-substrate adhesion. Orange and blue circles indicate vertices associated with cell-cell adhesion and cell-substrate adhesion, respectively. Large black circles indicate vertex 0. Crosses indicate cell centers. The following parameters were used: $K_t = 0.035$; $K_{adh} = 7.0 \times 10^{-7}$; $\eta = 3.0$.

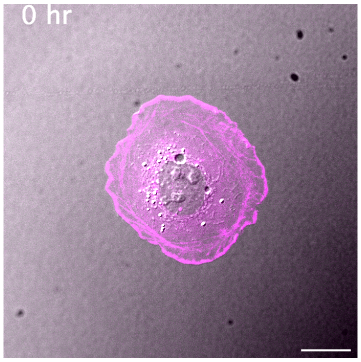

**Supplementary Video 11:** Clockwise rotation of a SHE78-7 treated single Caco-2 cell. DIC and Lifeact-RFP signals are shown in gray and magenta, respectively. Images were acquired every 15 minutes for 12 h. Scale bar, 20 $\mu$m. Display rate, 20 frames/sec.

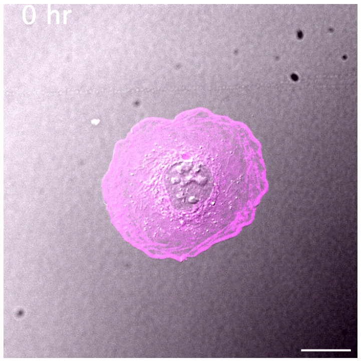

**Supplementary Video 12:** Clockwise rotation of a normal IgG treated single Caco-2 cell. DIC and Lifeact-RFP signals are shown in gray and magenta, respectively. Images were acquired every 15 minutes for 12 h. Scale bar, 20 $\mu$m. Display rate, 20 frames/sec.

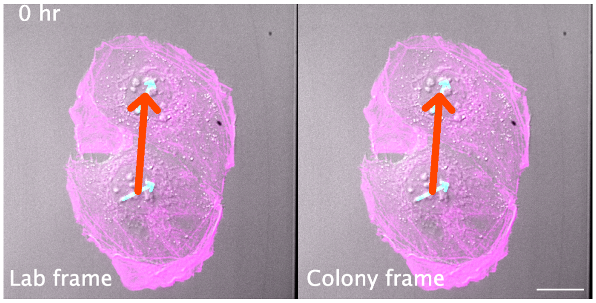

**Supplementary Video 13:** Counterclockwise collective rotation of a SHE78-7 treated two-cell Caco-2 colony. DIC and Lifeact-RFP signals are shown in gray and magenta, respectively. The left and right panels show motion in the laboratory frame and colony frame, respectively. Cyan and red arrows indicate vectors used to measure nuclear rotation and collective rotation. Images were acquired every 15 minutes for 12 h. Scale bar, 20 $\mu$m. Display rate, 20 frames/sec.

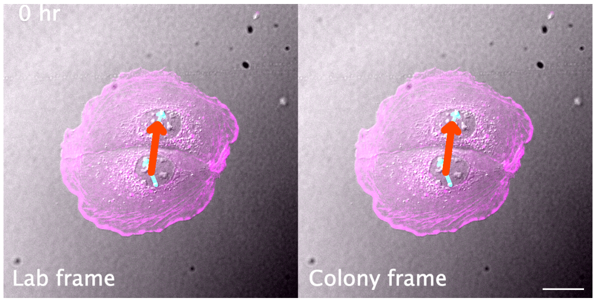

**Supplementary Video 14:** Clockwise collective rotation of a normal IgG treated two-cell Caco-2 colony. DIC and Lifeact-RFP signals are shown in gray and magenta, respectively. The left and right panels show motion in the laboratory frame and colony frame, respectively. Cyan and red arrows indicate vectors used to measure nuclear rotation and collective rotation. Images were acquired every 15 minutes for 12 h. Scale bar, 20 $\mu$m. Display rate, 20 frames/sec.

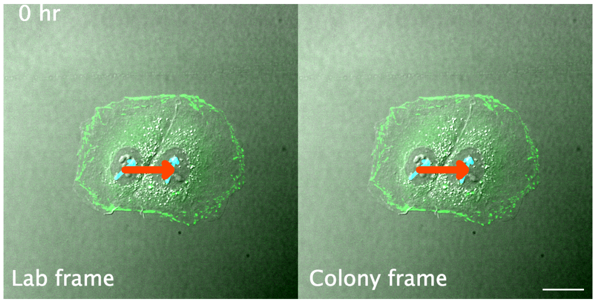

**Supplementary Video 15:** Slowed clockwise collective rotation of an FAK::GFP overexpressing two-cell Caco-2 colony. DIC and FAK::GFP signals are shown in gray and green, respectively. The left and right panels show motion in the laboratory frame and colony frame, respectively. Cyan and red arrows indicate vectors used to measure nuclear rotation and collective rotation. Images were acquired every 15 minutes for 12 h. Scale bar, 20 $\mu$m. Display rate, 20 frames/sec.

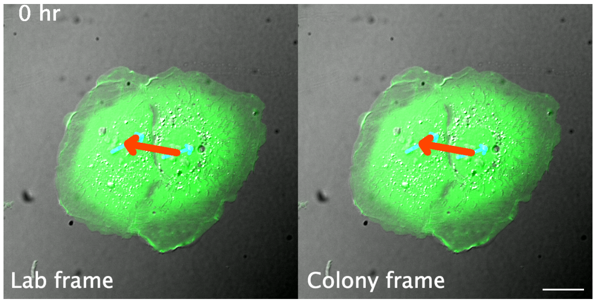

**Supplementary Video 16:** Clockwise collective rotation of an GFP overexpressing two-cell Caco-2 colony. DIC and GFP signals are shown in gray and green, respectively. The left and right panels show motion in the laboratory frame and colony frame, respectively. Cyan and red arrows indicate vectors used to measure nuclear rotation and collective rotation. Images were acquired every 15 minutes for 12 h. Scale bar, 20 $\mu$m. Display rate, 20 frames/sec.

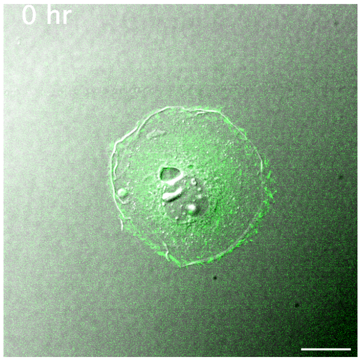

**Supplementary Video 17:** Slowed clockwise rotation of an FAK::GFP overexpressing single Caco-2 cell. DIC and FAK::GFP signals are shown in gray and green, respectively. Images were acquired every 15 minutes for 12 h. Scale bar, 20 $\mu$m. Display rate, 20 frames/sec.

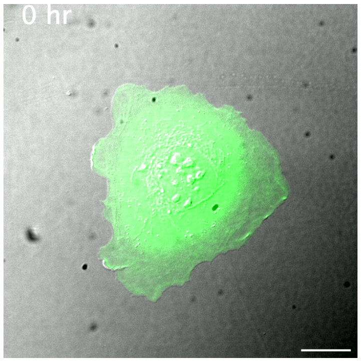

**Supplementary Video 18:** Clockwise rotation of an GFP overexpressing single Caco-2 cell. DIC and FAK::GFP signals are shown in gray and green, respectively. Images were acquired every 15 minutes for 12 h. Scale bar, 20 $\mu$m. Display rate, 20 frames/sec.

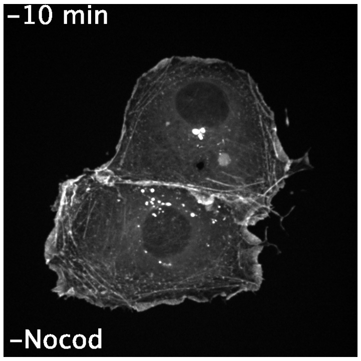

**Supplementary Video 19:** F-actin detachment from cell-cell junctions after nocodazole treatment. Lifeact-RFP signal is shown. The first frame shows the image before nocodazole treatment. Images were acquired every 10 min for 12 h after nocodazole treatment. Scale bar: 20 $\mu$m. Display rate is 20 frames/sec.

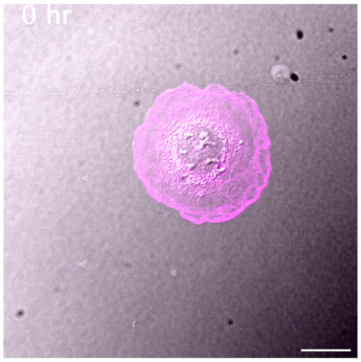

**Supplementary Video 20:** Clockwise rotation of an ACF7-siRNA treated single Caco-2 cell. DIC and Lifeact-RFP signals are shown in gray and magenta, respectively. Images were acquired every 15 minutes for 12 h. Scale bar, 20 $\mu$m. Display rate, 20 frames/sec.

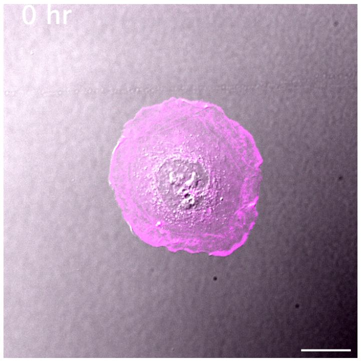

**Supplementary Video 21:** Clockwise rotation of an NC-siRNA treated single Caco-2 cell. DIC and Lifeact-RFP signals are shown in gray and magenta, respectively. Images were acquired every 15 minutes for 12 h. Scale bar, 20 $\mu$m. Display rate, 20 frames/sec.

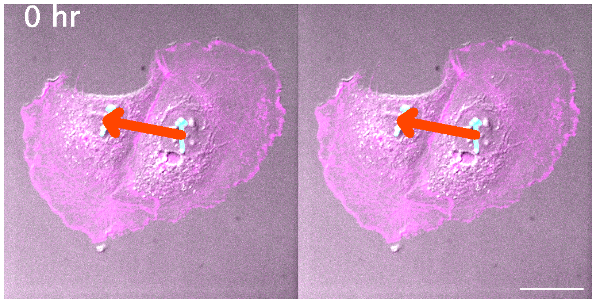

**Supplementary Video 22:** Counterclockwise collective rotation of an ACF7-siRNA treated two-cell Caco-2 colony. DIC and Lifeact-RFP signals are shown in gray and magenta, respectively. The left and right panels show motion in the laboratory frame and colony frame, respectively. Cyan and red arrows indicate vectors used to measure nuclear rotation and collective rotation. Images were acquired every 15 minutes for 12 h. Scale bar, 20 $\mu$m. Display rate, 20 frames/sec.

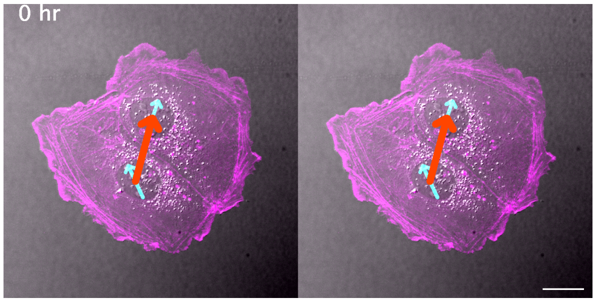

**Supplementary Video 23:** Clockwise collective rotation of an NC-siRNA treated two-cell Caco-2 colony. DIC and Lifeact-RFP signals are shown in gray and magenta, respectively. The left and right panels show motion in the laboratory frame and colony frame, respectively. Cyan and red arrows indicate vectors used to measure nuclear rotation and collective rotation. Images were acquired every 15 minutes for 12 h. Scale bar, 20 $\mu$m. Display rate, 20 frames/sec.

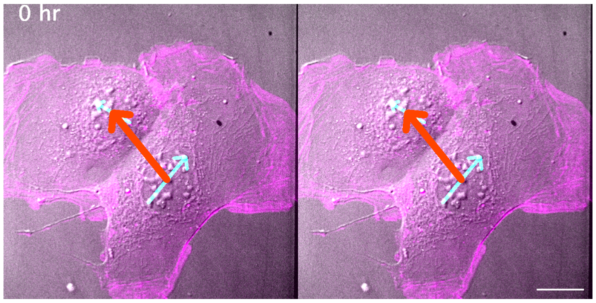

**Supplementary Video 24:** Counterclockwise collective rotation of an $\alpha$ECat-KO two-cell Caco-2 colony. DIC and Lifeact-RFP signals are shown in gray and magenta, respectively. The left and right panels show motion in the laboratory frame and colony frame, respectively. Cyan and red arrows indicate vectors used to measure nuclear rotation and collective rotation. Images were acquired every 15 minutes for 12 h. Scale bar, 20 $\mu$m. Display rate, 20 frames/sec.

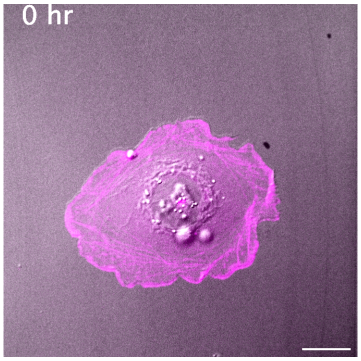

**Supplementary Video 25:** Clockwise rotation of an $\alpha$ECat-KO single Caco-2 cell. DIC and Lifeact-RFP signals are shown in gray and magenta, respectively. Images were acquired every 15 minutes for 12 h. Scale bar, 20 $\mu$m. Display rate, 20 frames/sec.

### Supplementary Notes

#### Theoretical model of the two-cell collective rotation

We describe here a theoretical description of the collective rotation of a two-cell colony of Caco-2 cells (Fig. 1). Based on our observations (Fig. 4; Extended Data Fig. 4), we hypothesized that the balance between cell-cell and cell-substrate adhesions determines the direction of the collective rotation. To test this hypothesis, we developed a theoretical description of the collective rotation of Caco-2 cells. In the model, the cell membrane and cell cortex require clockwise rotation, reflecting the single-cell chirality. In addition, both cell-cell and cell-substrate adhesions need to be included in the model. To satisfy these requirements, we employed a two-dimensional cell membrane model, where cells are described by polygons with vertices and edges. The cell chirality of individual cells is introduced as active forces working in the tangential direction along the cell periphery on vertices (Fig. 3a). The cell-cell adhesions work between the vertices of two cells (Fig. 3b). Finally, the cell-substrate adhesions are described by the frictional force exerted on the vertices. Reflecting the distribution of FAs (Supplementary Fig. 1c), the frictional coefficient is higher along the colony periphery than along the cell-cell interface.

Considering the force balance between the cell-substrate friction, potential forces, and active force, the equation of motion for the position $\vec{r}_i^\alpha$ of vertex $i$ $(i = 1, \cdots, N)$ of cell $\alpha$ ($\alpha$ =*I, II*) is given by

$$\eta_i^\alpha \frac{d\vec{r}_i^\alpha}{dt} = -\frac{\partial U}{\partial \vec{r}_i^\alpha} + \vec{T}_i^\alpha, \qquad (5)$$

where the first term on the left hand side is the frictional force and the first and second terms on the right hand side are potential forces and active force, respectively. The potential function $U$ in Eq. (5) is given by

$$U = U_a + U_l + U_b + U_{adh}, \qquad (6)$$

where $U_a$ is the area elasticity, $U_l$ is the line elasticity, $U_b$ is the bending rigidity, and $U_{adh}$ is the cell-cell adhesion. The area elasticity $U_a$ is the contribution from the hydrostatic pressure of cytoplasm and cytoskeleton, given by

$$U_a = \sum_\alpha \left[\frac{k_a}{2}(a^\alpha - a_0)^2\right], \qquad (7)$$

where $k_a$ is the area elastic modulus, $a^\alpha$ is the are of the $\alpha$-th cell, and $a_0$ is the preferred area of cells. The line elasticity $U_l$ originates from the tensile force exerted by the actin cortices along the cell membrane, given by

$$U_l = \sum_\alpha \sum_{i=0}^{N-1} \left[\frac{k_l}{2}\left(\frac{\ell_{i,i+1}^\alpha - \ell_0}{\ell_0}\right)^2\right], \qquad (8)$$

where $k_l$ is the line elastic modulus, $\ell_{i,i+1}^{\alpha}$ is the length of the segment connecting $i$ and $i+1$ vertices of cell $\alpha$, and $\ell_0$ is the preferred length of the segments. The bending rigidity $U_b$ of the membrane is given by

$$U_b = \sum_{\alpha} \sum_{i=0}^{N-1} [k_b(1 - \cos\theta_i^{\alpha})], \qquad (9)$$

where $k_b$ is the bending modulus and $\theta_i^{\alpha}$ is the angle of the two segments of the vertex $i$ in the cell $\alpha$. The cell-cell adhesion $U_{adh}$ is given by

$$U_{adh} = \sum_{i=0}^{N-1} \sum_{j=0}^{N-1} \left[k_{adh}\phi\left(d_{i,j}; d_0, d_{max}\right)\right], \qquad (10)$$

where $k_{adh}$ is the coefficient reflecting the level of the cell-cell adhesion and $\phi\left(d_{i,j}; d_0, d_{max}\right)$ is the product of the Morse potential $\psi_{Morse}\left(d_{i,j}\right)$ and the step function $s\left(d_{i,j}; d_0, d_{max}\right)$[38] given by

$$\phi\left(d_{i,j}; d_0, d_{max}\right) = \psi_{Morse}\left(d_{i,j}\right) s\left(d_{i,j}; d_0, d_{max}\right), \qquad (11)$$

where $d_{i,j}$ is the distance between vertex $i$ of the cell and vertex $j$ of the other cell, and $d_0$ and $d_{max}$ are paremeters. The Morse potential is given by

$$\psi_{Morse}\left(d_{i,j}\right) = \left[1 - \exp\left(\frac{d_0 - d_{i,j}}{w}\right)\right]^2 - 1, \qquad (12)$$

where $w$ is the coefficient that determines the width of the potential. The step function $s\left(d_{i,j}\right)$ is defined such that for $d_{i,j} \le d_0$, a repulsive force acts, for $d_0 < d_{i,j} < d_{max}$, an attractive force acts, and for $d_{i,j} \ge d_{max}$, there is no interaction on the vertex $i$ of the cell and vertex $j$ of the other cell. The step function $s\left(d_{i,j}\right)$ is given by

$$s\left(d_{i,j}; d_0, d_{max}\right) = \begin{cases} 1 & (d \le d_0) \\ \left(\frac{d_{max} - d}{d_{max} - d_0}\right)^3 \left[6\left(\frac{d_{max} - d}{d_{max} - d_0}\right)^2 - 15\left(\frac{d_{max} - d}{d_{max} - d_0}\right) + 10\right] & (d_0 < d < d_{max}) \\ 0 & (d \ge d_{max}). \end{cases} \qquad (13)$$

In the frictional force term on the left hand side of Eq. (5), $\eta_i^{\alpha}$ is the frictional coefficient of vertex $i$ of cell $\alpha$. The friction force reflects the frictional forces exerted by the cell-substrate adhesion. Here, the friction coefficient is $\eta_1$ for vertices where cell-cell adhesion is active ($d_{i,j} < d_{max}$), and $\eta_2$ for vertices where cell-cell adhesion is not active ($d_{i,j} \ge d_{max}$). Considering that focal adhesions involving cell-substrate adhesion are sparse beneath the cell boundaries where cell-cell adhesions are formed, we suppose

$$\eta_2 > \eta_1.$$

To describe the clockwise cell chirality observed in our Caco-2 cells, we introduced an active force to each vertex in the second term on the right hand side of Eq. (5). Since the active force originates from the cytoplasmic rotation motion, the force $\vec{T}_i^{\alpha}$ works in the tangential direction along the cell periphery, given by

$$\vec{T}_i^{\alpha} = \frac{k_t}{2}(\vec{r}_{i+1}^{\alpha} - \vec{r}_{i-1}^{\alpha}), \qquad (14)$$

where $k_t$ is the coefficient reflecting the level of the cell chirality.

To rescale time, length, and energy, we used the following units $\frac{\eta_1}{k_a p_0^2}$, $p_0$, and $k_a p_0^4$, respectively, where $p_0$ is the cell circumference. We obtained the non-dimensional equation of motion, and this equation contains the following non-dimensional parameters, cell-cell adhesion $K_{adh}$, friction coefficient $\eta$, active force $K_t$, preferred area $A_0$, segment elasticity $K_l$, bending rigidity $K_b$, equilibrium length of cell-cell adhesion $D_0$, maximum length of cell-cell adhesion $D_{max}$, and preferred length of segments $L_0$, as shown in Table 1. In the present paper, we solved this non-dimensional equation of motion with non-dimensional parameters.

We determine the value of $K_t$ of the control cells. $K_t$ is given by the dimensional parameters as

$$K_t = \frac{k_t}{\eta/\ell_0} \cdot \frac{\eta/\ell_0}{k_a p_0} \cdot \frac{1}{p_0}, \qquad (15)$$

where $\frac{k_t}{\eta/\ell_0}$ is the rotational velocity of individual cells, $\frac{\eta/\ell_0}{k_a p_0}$ is the relaxation time of cell area. Similarly, we estimated the cell circumference to be $\sim 50\pi$ $\mu$m. We also estimated the relaxation time of the cell area to be $\sim$ 0.25 hr, because cell rounding is resolved about 15 minutes after cell division.

#### Numerical simulation

For the numerical simulations, we used Euler's method with a constant time step $\Delta t = 0.001$. The initial positions of the vertices were set to the coordinates when the cell-cell adhesion reached equilibrium without exerting active force, followed by the calculation until $t = 100{,}000$. After discarding the initial transient ($t$ from 0 to 30,000), rotational velocities were calculated from the steady-state period ($t$ from 70,000 to 100,000). After the simulation, the final angles of the collective rotation $\Theta$ and individual cell rotations $\theta_{lab}$ were measured.

### Supplementary Table

**Table 1**: Non-dimensionalized parameters used in the theoretical simulations.

| | **Non-dimensionalized parameter** | **Definition** | **Non-dimensionalized value** |
|---|---|---|---|
| Cell-cell adhesion | $K_{adh}$ | $k_{adh}/k_a p_0^4$ | |
| Friction coefficient | $\eta$ | $\eta_2/\eta_1$ | |
| Active torque | $K_t$ | $k_t/k_a p_0^2$ | 0.035 |
| Preferred area | $A_0$ | $\pi R^2/p_0^2$ | ~ 0.08 |
| Segment elasticity | $K_l$ | $k_l/k_a p_0^4$ | ~ 50 |
| Bending rigidity | $K_b$ | $k_b/k_a p_0^4$ | ~ 0.01 |
| Equilibrium length of cell-cell adhesion | $D_0$ | $d_0/p_0$ | ~ 0.0038[39] |
| Maximum length of cell-cell adhesion | $D_{max}$ | $d_{max}/p_0$ | ~ 0.0076[40] |
| Preferred length of segments | $L_0$ | $\ell_0/p_0$ | ~ 0.005 |
| Number of vertices | $N$ | | 200 |
